# Low-dose radiopharmaceutical therapy enhances the efficacy of B7-H3 CAR T cells in murine metastatic neuroblastoma

**DOI:** 10.64898/2026.09.16.751528

**Authors:** Amanda G. Shea, Andrew Ollendorff, Andreanna Wilke, Lei Shi, Amy K. Erbe, Israrul H. Ansari, Malick Bio Idrissou, Ohyun Kwon, Hansel Comas Rojas, Marta Barisa, John Anderson, Jens C. Eickhoff, Irene Ong, Rebecca M. Richards, Paul M. Sondel, Christian M. Capitini, Reinier Hernandez, Bryan P. Bednarz, Jamey P. Weichert, Zachary S. Morris, Quaovi H. Sodji

## Abstract

**Background:** Chimeric antigen receptor (CAR) T cell therapy has had clinical success in hematologic malignancies, but limited efficacy is seen in solid tumors. In this study, we investigated whether systemic CAR T cell therapy could be enhanced in metastatic models of neuroblastoma when combined with radiopharmaceutical therapy (RPT).

**Methods:** Non-irradiated or irradiated tumor cells were co-cultured with CAR T cells (1:1) *in vitro* and supernatant media was subsequently collected for cytokines analyses. CAR T cells’ phenotypes were characterized by flow cytometry including checkpoint marker expression. Xenograft models of metastatic neuroblastoma were generated in NOD-*Rag1^null^IL2rg^null^*(NRG) mice. Tumor-bearing mice received 1.8 Gy of radiation delivered by ^177^Lu-NM600 RPT five days after tumor implantation. Nine days after RPT, 4 x 10^6^ CAR T cells were administered intravenously. To evaluate tumor burden, mice were imaged weekly for 4 weeks.

**Results:** In models of metastatic neuroblastoma, ^177^Lu-NM600 RPT significantly increased overall survival when combined with CAR T cell therapy *in vivo*. Pre-treatment of tumor cells with ^177^Lu also significantly increased CAR T cell cytotoxicity while decreasing production of IL-4 and IL-10 *in vitro*. Co-culture of CAR T cells with irradiated tumors led to increases in PD-1^+^TIM3^+^LAG3^+^ T cells, suggesting that further combination with immune checkpoint inhibitors may enhance clinical efficacy.

**Conclusions:** Our findings demonstrate that low-dose RPT can potentiate the anti-tumor efficacy of CAR T cells in metastatic neuroblastoma. To our knowledge, this is the first report of dosimetry-based RPT being combined with CAR T cells in a metastatic solid tumor setting. These findings underscore the potential of combining RPT and CAR T cells to overcome the unique challenges of solid tumors, particularly when treating metastatic disease.

## Introduction

Immunotherapy has fundamentally changed the treatment landscape for many cancers. This includes adoptive cell therapies (ACT) (1, 2) such as chimeric antigen receptor T cells (CAR T), a novel immunotherapy that has been FDA approved for the treatment of hematological malignancies including non-Hodgkin lymphoma (3), multiple myeloma (4) and B cell lymphoblastic leukemia (5). CAR T cell treatment in these contexts has been shown to be generally safe and effective, with overall response rates ranging from 40-90%. This suggests great promise for the application of CAR T cells in the treatment of other cancers. However, the success in hematologic malignancies has not been uniformly replicated in solid tumors. Poor CAR T cell trafficking and limited infiltration into the immunosuppressive tumor microenvironment (TME), T cell exhaustion, and tumor antigen escape have all contributed to the limited clinical efficacy of CAR T cells in solid tumors (6).

To mitigate the poor responses of CAR T cells in solid tumors, studies have begun combining radiation with CAR T cells. Radiotherapy, such as delivered by external beam radiation (EBRT), facilitates immunogenic cell death (7), which has been shown to promote immune activation, priming, and T cell migration into the TME (8). Emerging pre-clinical data suggests that in localized disease, EBRT synergizes with CAR T cells to enhance anti-tumor efficacy and increase persistence (9, 10). While a promising combination treatment modality, delivering EBRT to radiographically occult and metastatic lesions would generally require whole-body radiation, which can induce severe toxicities including lymphopenia and immunosuppression (11, 12).

Radiopharmaceutical therapy (RPT) is a form of systemically administered radiation therapy that uses a tumor targeting moiety to selectively deliver a radioactive isotope to tumors following intravenous administration (13). RPT can selectively deliver radiation to tumors even in metastatic settings while minimizing radiation damage to healthy tissues and lowering the potential for toxicity compared to whole-body EBRT (13). Due to variations in perfusion and tumor target cell surface distribution (14), RPT delivers a heterogeneous radiation dose to tumors. Emerging evidence suggests that by providing a low, moderate, and high radiation dose to a single tumor microenvironment (TME), RPT may engage multiple immune modulatory pathways (15). RPT generates anti-tumor responses when combined with other immunotherapies, such as immune checkpoint blockade (16) and CAR T cells in non-metastatic settings (17), but it is not yet known whether RPT can enhance the efficacy of CAR T cells in the metastatic setting.

Amongst RPT targeting moieties currently under investigation, NM600, an alkylphosphocholine analog, preferentially accumulates in the membranes of most solid tumor cells (18, 19). Here, we evaluated the ability of low-dose RPT delivered by [177Lu]Lu-NM600, NM600 conjugated to the β-emitter Lutetium-177, (^177^Lu-NM600) to enhance CAR T cell therapy efficacy against metastatic neuroblastoma xenograft models.

## Materials and Methods

### Cell lines

The human neuroblastoma cell lines CHLA-20 (Children’s Oncology Group), SH-SY5Y (ATCC), and SK-N-AS (ATCC) were used. CHLA-20-AkaLUC-GFP was generously provided by Dr. James Thomson (University of Wisconsin-Madison). SH-SY5Y-LUC-GFP was generously provided by Dr. Crystal Mackall (Stanford University). SK-N-AS was provided by Dr. John Maris (Children’s Hospital of Philadelphia). Cells were cultured in Dulbecco’s Modified Eagle Medium (DMEM) (Gibco, #10-013-CV) with high glucose, supplemented with 10% fetal bovine serum (FBS)(Avantor) and 1% penicillin-streptomycin (P/S) (Gibco, #15140163), and maintained at 37°C in a 5% CO_2_ atmosphere and used after 3-5 passages in culture after thawing. Cell authentication was performed per ATCC guidelines using morphology, growth curves, and Mycoplasma testing within 6 months of use with the MycoStrip Mycoplasma Detection Kit (Invitrogen, Waltham, MA).

### Human T cell isolation and CAR T cell manufacturing

Primary human T cells were isolated from leukocyte reduction system (LRS) cones (Versiti) using negative selection with the EasySep Human T Cell Isolation Kit (Stemcell Technologies, #100-0695) or were obtained from healthy donor Leukopaks (Versiti) that were previously TCRαβ-positively sorted and kindly donated by Dr. Jacques Galipeau (University of Wisconsin-Madison). Primary T cell phenotype was confirmed by flow cytometry.

The second-generation B7-H3-CD28-CD3ζ CAR construct was generated in the γ-retroviral producer line Pheonix-ampho-HEK293T^21^. ImmunoCult-XF media (Stemcell Technologies, #10981) was supplemented with 1% P/S. Isolated T cells were stimulated by either anti-CD3 (Invitrogen, #16-0037-85 and anti-CD28 (Invitrogen, #14-0289-82) as soluble antibodies or as Dynabeads (Gibco, #11132D) and cultured in ImmunoCult-XF media supplemented with 1% P/S and 100 IU/mL IL-2 (Tecnin). After 72 hours of activation, viral transduction retronectin (Takara, #T100B) plates were prepared by spinoculation (2500xg, 120 min, 32°C). Activated T cells were transfected for 48 hours on transduction plates. Anti-CD3 and anti-CD28 antibodies were then removed, and CAR T cells were expanded in ImmunoCult-XF media supplemented with 1% P/S and 100 IU/mL IL-2 until day 12. CAR T cells were assessed for CAR expression by flow cytometry and cryopreserved in Cryostor CS10 (Biolife Solutions, #210102). Cryopreserved CAR T cells were thawed and resuspended in ImmunoCult-XF media supplemented with 1% P/S and 50 IU/mL IL-2 for 24 hours at 37°C, 5% CO_2_ prior to *in vitro* or *in vivo* use.

### *In vitro* dosimetry, *In vivo* dosimetry, and Radiosynthesis of ^177^Lu-NM600

The specific activities of ^177^Lu required to deliver 2 Gy to a cell monolayer at specific time points were estimated using the Geant4 Monte Carlo toolkit with an extension of RAPID, as previously described (19, 20). *In vivo* dosimetry of ^177^Lu-NM600 was estimated as previously reported (17, 20, 21). The total activity in each organ was calculated by extrapolating %ID/g (percent injected dose per gram) at given time points (Supplemental Figure S1A-C). The radiolabeling of NM600 with ^177^LuCl_3_ (SHINE Technologies) was performed as previously reported (21).

### Flow cytometry

Tumor cells or CAR T cells were resuspended into FACS buffer and stained with Fixable Live Dead-NIR (ThermoFisher, #L34994) (20 min, RT, dark). Surface staining was performed using fluorophore-conjugated antibodies (30 min, RT, dark). Flow cytometry antibodies are listed in Supplementary Table S1. Compensation was performed using UltraComp eBeads (ThermoFisher, #01-2222-42) or single stained cells. Data were acquired on the Attune (ThermoFisher) and analyzed with either R (Posit) or FlowJo (TreeStar). Following flow cytometry analysis, Pearson residuals analysis of the multi-checkpoint status of CAR T cells following co-culture with either non-irradiated or irradiated tumor cells was assessed. Magnitude of the Pearson residual (degree of deviation between the expected vs observed counts), directional significance (signed –log10(p)), or differentiation of a specific phenotype relative to the total population was determined. Following flow cytometry analysis of median fluorescence intensity (MFI), differences in B7-H3 expression (log_10_MFI) across cell lines were analyzed using a linear mixed-effects model. The cell line was treated as a fixed effect, while the experiment was included as a random effect to account for inter-experimental variation.

### Flow cytometric analysis of neuroblastoma cells following RPT treatment *in vitro*

Each neuroblastoma cell line was incubated with the activity of ^177^Lu to achieve 2 Gy over 3 days. After 72 hours, viability of non-irradiated or irradiated neuroblastoma cells were assessed by flow cytometry. Biological replicates were N = 3, and technical replicates were n = 3. Differences in B7-H3, HLA-DR, Galectin 9, CD155, and PD-L1 expression or MFI across cell lines were analyzed using a linear mixed-effects model. Radiation dose (0 Gy vs 2 Gy) was treated as a fixed effect, while experimental batch (biological replicate) was included as a random effect to account for the nested structure of technical replicates.

### *In vitro* cytotoxicity activity of B7-H3 CAR T cells

#### Non-radiation Experiments

CHLA-20-AkaLUC-GFP, SH-SY5Y-LUC-GFP, and SK-N-AS-mKate neuroblastoma cell lines were plated for cytotoxicity assays at 1 x 10^4^ cells per well. Thawed B7H3 CAR T cells were co-cultured at an effector-to-target (E:T) ratio of 4:1, 2:1, or 1:1 for 24 hours by the Incucyte S3 (Sartorius). Images were taken every 4 hours for 24 hours. Green Integrated Intensity per image or Red Integrated Intensity per image was used to calculate the relative intensity of GFP+ (CHLA-20 and SH-SY5Y) or mKate+ (SK-N-AS) neuroblastoma cells in each well and normalized to the first image taken. Images were analyzed using the IncuCyte Analysis Software (Sartorius). Each experiment was repeated across two donors.

#### Radiation Experiments

Neuroblastoma cell lines were incubated with the activity of ^177^Lu to achieve 2 Gy over 3 days. After 72 hours, viability of non-irradiated or irradiated neuroblastoma cells were assessed by flow cytometry or plated for a cytotoxicity assay. Tumor cells were washed several times with DMEM with high glucose, supplemented with 10% FBS and 1% P/S at 37 °C and 5% CO_2_. Thawed CAR T cells were co-cultured with either non-irradiated or irradiated tumor cells at an E:T ratio of 1:1 for 24 hours by Incucyte S3 (Sartorius). Images were taken every 4 hours for 24 hours. Green Integrated Intensity per image or Red Integrated Intensity per image was used to calculate the relative intensity of GFP^+^ or mKate^+^ neuroblastoma cells in each well and normalized to the first image taken. Images were analyzed using the IncuCyte Analysis Software (Sartorius). The area under the curve (AUC) was calculated for each experiment. Biological replicates for were N = 3 and technical replicates were n = 3. Statistical significance was determined using a linear mixed model with treatment (cell line) and radiation as fixed effects and the experiments as random effects to account for the nested structure of the technical replicates. Degrees of freedom were estimated using the Kenward-Roger approximation.

### *In vitro* cytokine detection assays

Twenty-four hours after co-culturing CAR T cells with non-irradiated or irradiated neuroblastoma cells, supernatant was collected and stored at -80°C until supernatant reached background radioactivity. The levels of selected cytokines were assessed using the LEGENDplex Human CD8/NK Panel V02 (Biolegend) according to the manufacturer’s protocol and collected on the Attune (ThermoFisher). Data were analyzed with the Qognit (Biolegend) software to determine the pg/mL concentration. Biological replicates were N = 2, and technical replicates were n = 3. Statistical significance was determined using a linear mixed model with radiation dose as a fixed factor and the biological experiments as a random effect to account for the nested technical replicates.

### *In vivo* human xenograft neuroblastoma model treated with ^177^Lu-NM600 RPT and B7-H3 CAR T cells

All animal experiments were approved by the University of Wisconsin-Madison Animal Care and Use Committee (IACUC protocol M006815). 9-12 week old male or female NOD-*Rag1^null^IL2rg^null^* (NRG) mice (Jackson Laboratory) were injected intravenously by tail vein with 0.5 x 10^6^ CHLA-20-AkaLUC-GFP or SH-SY5Y-LUC-GFP to establish metastatic neuroblastoma tumors. Tumor engraftment was verified using bioluminescence measurements on the Lago In Vivo Imaging System (Spectral Instruments Imaging). To evaluate bioluminescent imaging (BLI), mice were sedated with isoflurane and given either AkaLumine (CHLA-20, 1.25mM, 100 μL) (MediLumine, #305AKA) or 100 μL/mouse of d-Luciferin (SH-SY5Y, 30 mg/mL, 100 μL) (MediLumine, #222PS) by intraperitoneal injection. Lago images were taken prior to CAR T injection to establish pre-CAR tumor sizes. Mice were then randomized into the following groups: No Treatment, CAR T cells only, ^177^Lu-NM600 alone, ^177^Lu-NM600 + untransduced (UTD) T cells, and ^177^Lu-NM600 + CAR T cells. Groups contained 4-5 mice per group. On day 5 post-engraftment, ^177^Lu-NM600 RPT was administered by tail vein injection to CHLA-20 (50 μCi) or SH-SY5Y (80 μCi), to achieve 1.8 Gy.

On day 14, 4×10^6^ CAR^+^ T cells, as determined by flow cytometry, were administered via the tail vein. Mice were imaged weekly for BLI. To quantify tumor burden, a region of interest (ROI) was drawn around established tumors and total flux (photons/sec) was calculated by Aura In Vivo Imaging Software (Spectral Instruments Imaging). To evaluate tumor burden, the AUC was calculated from tumor BLI normalized to the first image (day 7 post-RPT, but prior to CAR administration). AUC was assessed for each mouse, and values were normalized to *log*_10_. Weights were recorded in grams and normalized to the day 7 weight.

### *In vivo* CAR T cell tracking

9 -12 week old male or female NRG mice (Jackson Laboratory) were injected intravenously by tail vein with 1 x 10^6^ CHLA-20-AkaLUC-GFP or SH-SY5Y-LUC-GFP cells to establish metastatic neuroblastoma tumors. Tumor engraftment was verified using BLI measurements on the Lago In Vivo Imaging System. To evaluate BLI, mice were sedated with isoflurane and were given either AkaLumine (CHLA-20, 1.25 mM, 100 μL) (MediLumine, #305AKA,) or 100 μL/mouse of d-Luciferin (SH-SY5Y, 30 mg/mL, 100 μL) (MediLumine, #222PS) by intraperitoneal injection. Mice were randomized into the following groups: No Treatment, B7H3 CAR T cells alone, or ^177^Lu-NM600 + B7H3 CAR T cells. CHLA-20 groups had n = 3 in the no-treatment group and n = 4 in the CAR T groups. SH-SY5Y had 5 mice per group. All mice were followed weekly for BLI measurements, weight loss, and overall survival.

On day 5 post-tumor engraftment, ^177^Lu-NM600 was administered by tail vein injection to CHLA-20- (50 μCi) or SH-SY5Y- (80 μCi) bearing mice to achieve 1.8 Gy. BLI images were taken prior to CAR T cell thaw to establish pre-CAR tumor sizes. On day 14, CAR T cells were labeled with IVISense DiR 750 (Revvity, #125964) fluorescent cell labeling dye for 30 minutes according to the manufacturer’s instructions. 4 x 10^6^ CAR^+^ T cells were administered intravenously by tail vein. Mice were BLI imaged 3, 24, 48, 72, and 240 hours after CAR T cell administration. To quantify T cells, fluorescent imaging (FLI) was performed with excitation and emission at 710/760 nm. A region of interest was drawn around established tumors to determine the tumor BLI and the T cell FLI. T cell trafficking was quantified by Aura In Vivo Imaging Software (Spectral Instruments Imaging). CAR T cell FLI trafficking data were analyzed by AUC.

### Flow cytometric analysis of *in vivo* CAR T cell homing and persistence

9–12-week-old female NRG mice (Jackson Laboratory) were intravenously injected by tail-vein with 0.5×10^6^ SH-SY5Y-LUC-GFP cells to establish metastatic tumors, and tumor engraftment was verified by BLI on the Lago In Vivo Imaging System. The mice were randomized into the following groups: No Treatment, B7H3 CAR T cells only, or B7H3 CAR T cells + ^177^Lu-NM600 RPT, with 14 mice per group. On day 5 post-engraftment, ^177^Lu-NM600 RPT was administered by tail vein injection to SH-SY5Y (80 μCi), to achieve 1.8 Gy in the radiation groups. On day 14, 4×10^6^ CAR^+^ T cells were injected into the CAR T groups intravenously by tail vein. Both 4 and 12 days after CAR T administration, the first cohort (n = 7) was sacrificed for isolation of the liver and spleen, respectively. Half of the liver and spleen was fixed in 10% formalin for immunohistochemistry (IHC). The other halves were disaggregated, filtered (40 μM), RBC lysed (Biolegend, 420301), and processed into single cell suspensions for analysis by flow cytometry.

### Immunohistochemistry of CAR T cells in mouse liver and spleen

Harvested spleen and liver were fixed in 10% formalin for 24 hours and stored in 70% EtOH (4°C). After reaching background ^177^Lu radioactivity level (70 days) tissue samples were paraffin-embedded (n = 3 from each group). Slides were generated from the formalin-fixed paraffin embedded samples. Hematoxylin and eosin (H&E) staining was performed. Samples were then stained for human CD45 (Cell Signaling, 13917T) (overnight, 4°C) to measure CAR T cell homing. The following day, samples were washed and secondary stained with DAB. Images were taken at 10X on the Evos (Invitrogen). 3 fields per sample were counted for CD45^+^ cells and averaged.

### *In vivo* CAR T cell treatment of human SK-N-AS neuroblastoma xenografts

9-12-week-old male or female NRG mice (Jackson Laboratory) were tail-vein injected with 1 × 10^6^ SK-N-AS-LUC cells to establish metastatic neuroblastoma tumors. On day 5, mice were randomized into the following groups: No Treatment (n = 5) or B7H3 CAR T cells (n = 5). On day 12 post-tumor engraftment, CAR T cells were thawed according to the CAR T cell thaw protocol as above. On day 13, 4 x 10^6^ CAR^+^ T cells were intravenously injected by tail vein. Mice were imaged on the IVIS weekly after being sedated with isoflurane and intraperitoneal injections of d-Luciferin (MediLumine, #222PS, 30mg/mL, 100 μL). All mice were followed for weight loss and overall survival.

### Data analysis and software

All data and statistical analyses, and graph generation were performed in GraphPad Prism (v.10.0.2), Microsoft Excel, or R (Version 2026.01.0+392). FlowJo (10.10.0, TreeStar) was used to analyze .fcs files exported from Attune NxT software (ThermoFisher). Figures were organized using Microsoft PowerPoint. Schemas within figures were generated by BioRender. R code generation was assisted by Google Gemini. R libraries can be found in Supplemental Table S2 (22–36). Post-hoc pairwise comparisons were performed using Tukey’s Honest Significant Difference (HSD) test to identify specific differences in B7-H3 MFI between tumor cell lines. For *in vitro* flow data, a one-way ANOVA was used to determine significance. For CAR T cytotoxicity data, planned post-hoc comparisons were performed using the estimated marginal means with Sidak’s adjustment for multiple testing. For *in vivo* studies, tumor burden was analyzed by one-way ANOVA followed by planned comparisons using the Holm-Sidak adjustment, comparing the combination therapy group CAR T + ^177^Lu-NM600 to all other experimental cohorts. The probability of survival was calculated using the Mantel-Cox Log-Rank test, where CAR T + ^177^Lu-NM600 was set as the reference group and compared to all other groups. CAR T cell FLI trafficking data were analyzed by Dunnett’s T3 multiple comparisons test. Homing flow data was analyzed using a two-way ANOVA with uncorrected Fisher’s LSD. A Welch’s t-test was used to determine significance on IHC images.

## Results

### Virally generated B7-H3 CAR T cells exhibit potent anti-tumor efficacy against target models of neuroblastoma *in vitro* and *in vivo*

We produced and expanded second-generation B7-H3 CAR T cells from three healthy donors (Figure 1A-B) then assessed CAR expression (Figure 1C, Supplemental Figure S2). As expected, CAR expression varied amongst the donors, averaging 60%. Thawed untransduced and CAR T cells from these donors revealed differing proportions of CD4 and CD8 cells (Figure 1D, Supplemental Figure S3A) but similar levels of exhaustion markers between donors after *ex vivo* expansion with CD3/CD28 and IL-2 (Supplemental Figure S3A-B).

**Figure 1.**
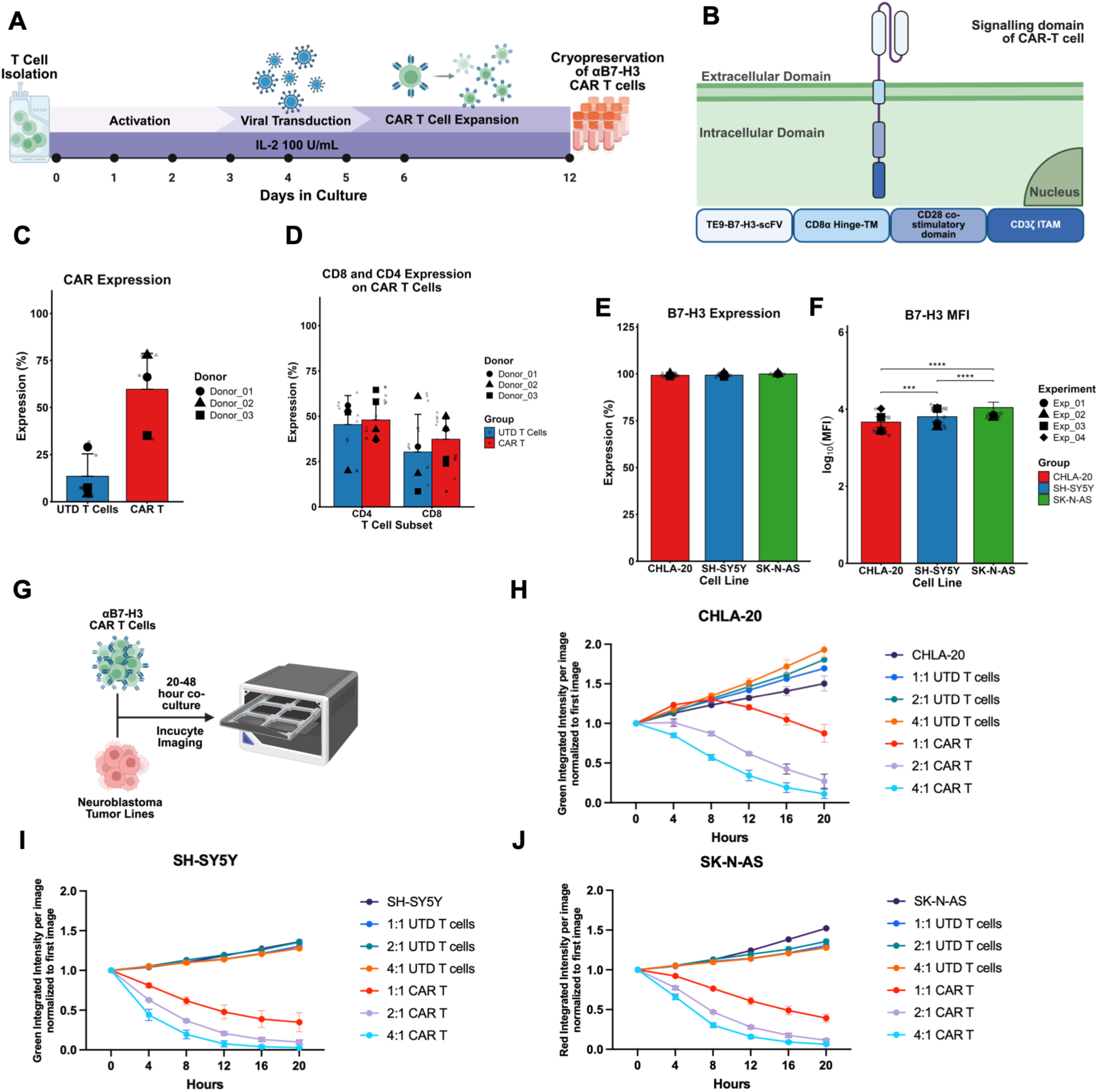
αB7-H3 CAR Ts have potent cytotoxicity against multiple models of neuroblastoma. **dA-B.** Schematic of αB7-H3 CAR T cell manufacture and representation of the αB7-H3 CAR T construct. αβ-T cells were isolated from leukopaks and stimulated with either CD3/CD28 soluble antibodies and IL-2 or Dynabeads and IL-2. On day 3, viral supernatant containing αB7-H3 CAR was inoculated with the T cells. Figures were created in BioRender. **C-D.** Flow analysis of **(C)** Expression of αB7-H3 CAR on T cells in three donors before cryopreservation. **(D)** of post-thaw CD8 and CD4 expression on 3 donors. Data are displayed as a superplot bar graph with large symbols representing the mean of three technical replicates, each represented by small, transparent symbols. Circle: Donor 1, square: Donor 2, and triangle: Donor 3. Error bars represent the ± SD of the biological means. **E-F.** Flow analysis of CHLA-20, SH-SY5Y, and SK-N-AS **(E)** percent expression and **(F)** median fluorescence intensity (MFI) of CAR T target antigen B7-H3. Differences in B7-H3 expression (log_10_MFI) were analyzed using a linear mixed-effects model. Data are represented in a superplot bar graph with N = 4 biological experiments (large symbols), each performed with n = 3 technical replicates (small, transparent symbols). Error bars represent the ± SE of the biological means. Statistical significance was determined by Tukey-adjusted pairwise comparison. **G.** Schematic of *in vitro* cytotoxicity assay. Untransduced T cells (UTD) and CAR T cells were harvested and plated at either 4:1, 2:1, or 1:1 effector to target (E:T) ratios against each of the tumor lines. **H-J.** Donor 2 cytotoxicity targeting CHLA-20, SH-SY5Y, and SK-N-AS as measured by decrease in green fluorescence intensity normalized to the first image. Significance is indicated as: ns (not significant), *\* (p < 0.05), ** (p < 0.01), *** (p < 0.001). ****(p < 0.0001)*.

We assessed B7-H3 expression in the CHLA-20, SH-SY5Y, and SK-N-AS neuroblastoma cell lines. While 100% of the cells in each of the cell lines expressed B7-H3, the MFI varied across tumor cell lines (Figure 1E and 1F, Supplemental Figure S4A-B). B7-H3 expression was higher in SK-N-AS than in either CHLA-20 or SH-SY5Y (Figure 1F). We next determined the cytotoxicity of B7-H3 CAR T cells against neuroblastoma cell lines *in vitro*, demonstrating killing at a low effector to target (E:T) ratio of 1:1 by donor 2 (Figure 1G-J) and donor 3 CAR T cells (Supplemental Figure S4C-D). To confirm the potency of our manufactured B7-H3 CAR T cells *in vivo*, we administered B7-H3 CAR T cells in a metastatic SK-N-AS neuroblastoma xenograft model. The probability of survival was significantly increased following treatment with CAR T cells compared with no treatment alone (Supplemental Figure S5A-C). Together, these data show that our manufactured B7-H3 CAR T cells have anti-tumor activity against three distinct neuroblastoma cell lines and in a metastatic neuroblastoma xenograft model when given as monotherapy.

### Low-dose RPT enhances B7-H3 CAR T cytotoxicity *in vitro*

Given that B7-H3 CAR T cells have been shown to be antigen density dependent (37, 38) and that radiation at moderate doses can increase the density of B7-H3 on certain tumor types (39), we assessed B7-H3 expression on neuroblastoma cells (CHLA-20, SH-SY5Y, and SK-N-AS) following low-dose radiation delivered by treatment of neuroblastoma with ^177^Lu (Figure 2A). B7-H3 MFI decreased on irradiated CHLA-20 cells (Figure 2B) despite no change in the percentage of CHLA-20 cells expressing B7-H3 (Supplemental Figure S6A). In contrast, both MFI and the percent of cells expressing B7-H3 were unchanged by ^177^Lu for SH-SY5Y (Figure 2C and Supplemental Figure S6B) and SK-N-AS cells (Supplemental Figure S6C-D). These data suggest that B7-H3 expression is not consistently altered by low-dose ^177^Lu in neuroblastoma cell lines.

**Figure 2.**
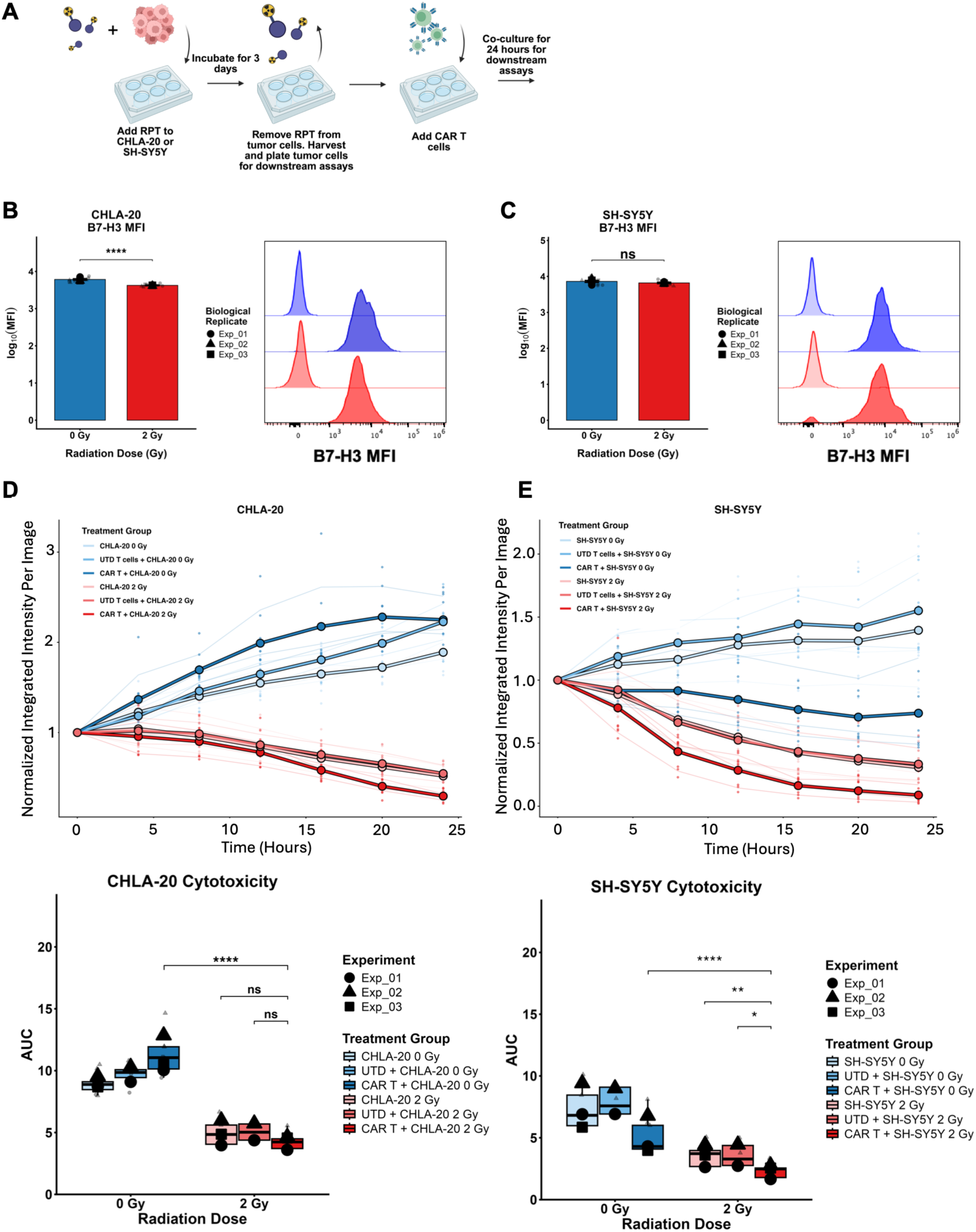
Low dose RPT enhances *in vitro* cytotoxicity. **A.** Schema of the experimental design. **B-C.** B7-H3 percent expression and median fluorescence intensity (MFI) by flow cytometry in (B) CHLA-20 and (C) SH-SY5Y. Data are represented as N = 3 biological experiments (large symbols), each performed with n = 3 technical replicates (small, transparent symbols). Error bars represent ± SE of the biological means. The histogram is a representative of one sample, while the initial peak corresponds an isotype. Statistical analysis of surface marker expression was performed using a linear mixed model (LMM), and significance was determined by a one-way ANOVA. **D-E.** Cytotoxicity assay after the indicated treatments to **(D)** CHLA-20 and **(E)** SH-SY5Y. αB7-H3 CAR T and UTD T cells were added in a 1:1 E:T ratio. Green integrated intensity per image was normalized to the first time point. Cytotoxicity time course is represented as line graphs of the grand mean of n = 3 biological replicates and n = 3 technical replicates. Area under the curve (AUC) data are presented as superplot boxplots showing three independent biological experiments (N = 3) and technical replicates (n = 3). Statistical significance was determined using a linear mixed model with planned estimated margins of means with Sidak’s adjustment for multiple testing. Significance is indicated as: ns (not significant), *\* (p < 0.05), ** (p < 0.01), *** (p < 0.001). ****(p < 0.0001)*.

We next determined if low-dose radiation delivered by ^177^Lu enhanced tumor cell susceptibility to CAR T cell cytotoxicity *in vitro*. We co-cultured B7-H3 CAR T cells at a 1:1 E:T ratio with either non-irradiated or irradiated tumor cells with 2 Gy free ^177^Lu. Tumor cell death increased significantly when CAR T cells were co-cultured with irradiated CHLA-20 tumor cells compared to non-irradiated CHLA-20 cells (Figure 2D). In contrast to CHLA-20, SH-SY5Y demonstrated significantly increased cell death across all non-irradiated treatment groups, as well as with ^177^Lu alone or in combination with untransduced (UTD) T cells (Figure 2E). Similar results were observed with donor 3 CAR T cells (Supplemental Figure S6E-F) and donor 2 CAR T cells cultured with SK-N-AS (Supplemental Figure S6G).

These data suggest that low-dose radiation may potentiate B7-H3 CAR T cell-mediated killing of neuroblastoma, but this effect may depend on B7-H3 antigen density and may be cell line dependent.

### Enhanced efficacy of B7-H3 CAR T cells co-administered with low-dose RPT in metastatic neuroblastoma

Our group has recently shown that low-dose radiation by RPT could render mice bearing a single neuroblastoma tumor implanted in the flank tumor-free when combined with GD2 CAR T cells^18^. Given differences in metastatic disease versus primary or localized tumors, and the potential clinical opportunity of combining RPT with CAR T cells, we examined the efficacy of ^177^Lu-NM600 RPT with B7-H3 CAR T cells *in vivo* using a xenograft model of metastatic neuroblastoma. Prior work showed that the residual radioactivity within the TME following 3.6 Gy of ^177^Lu-NM600 was deleterious to the viability of CAR T cells compared to the lower dose of 1.8 Gy (17). We therefore evaluated 1.8 Gy ^177^Lu-NM600 in NRG mice bearing human neuroblastoma metastatic to the liver.

After confirming metastatic tumor engraftment by bioluminescence (BLI), mice were randomized to receive no treatment, B7-H3 CAR T cells alone, ^177^Lu-NM600 (1.8 Gy) alone, UTD T cells + ^177^Lu-NM600 (1.8 Gy), or B7-H3 CAR T cells + ^177^Lu-NM600 (1.8 Gy) (Figure 3A). Each tumor-bearing RPT-treated mouse received enough activity of ^177^Lu-NM600 to achieve a mean absorbed tumor dose of 1.8 Gy, corresponding to 50 μCi for CHLA-20 (Figure 3B-E) or 80 μCi for SH-SY5Y (Figure 3F-I). Nine days post-RPT treatment, the mice were injected with either UTD T cells or B7-H3 CAR T cells and were imaged for BLI weekly until 4 weeks after RPT administration to assess tumor burden (Figure 3B and 3F).

**Figure 3.**
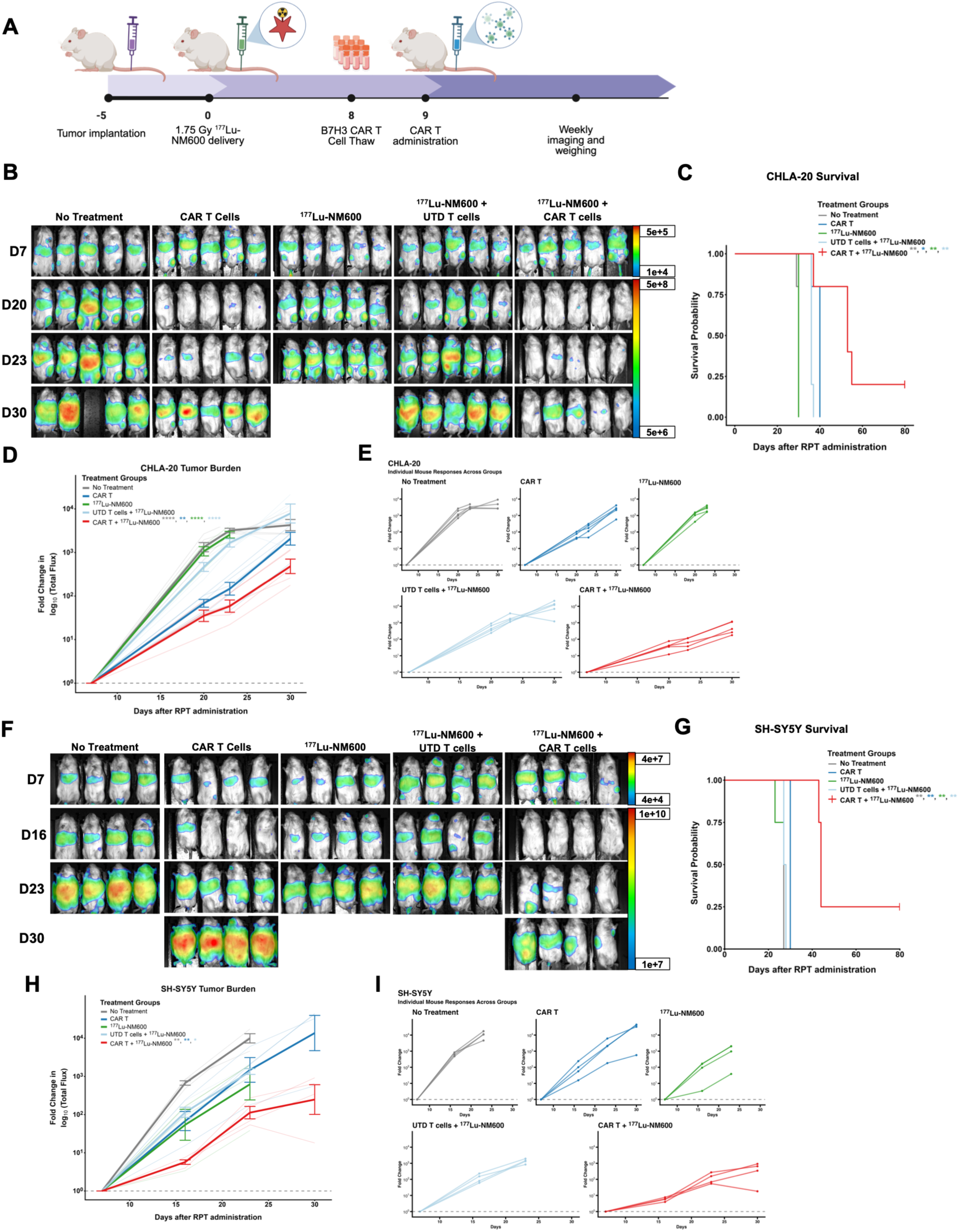
CAR T cell therapy efficacy is enhanced by low dose RPT pre-treatment to models of metastatic neuroblastoma. **A.** Schema of experiment timeline. **B-I.** Tumors were analyzed in mice bearing either **(B-E)** CHLA-20 or **(F-I)** SH-SY5Y after being treated as in (A). **(B, F)** Representative bioluminescence (radiance: photons/sec per square centimeter per steradian) Lago images of all treatment groups in the metastatic CHLA-20 tumor model. The first row of images (D7) represents the mice after RPT but before CAR T cell administration. (**C, G)** Kaplan-Meier curves illustrating the survival probability. Statistical significance was determined using the Mantel-Cox Log-Rank test. Average **(D, E)** and individual **(H, I)** tumor burden curves normalized to day 7. Area under the curve (AUC) was measured for each mouse, and values were normalized to log10 transformed, then analyzed by a one-way ANOVA followed by Holm-Sidak adjustment with CAR T + ^177^Lu-NM600 as the reference group. Metastatic tumor burden represented over time in individual mice in each treatment group. Significance is indicated as: ns (not significant), *\* (p < 0.05), ** (p < 0.01), *** (p < 0.001). ****(p < 0.0001)*.

In CHLA-20-bearing mice, tumor burden measured by BLI was significantly reduced in the combination RPT and B7-H3 CAR T cell group compared to all other treatment conditions (Figure 3D-E). Importantly, in both the CHLA-20 and SH-SY5Y metastatic models, the combination RPT and B7-H3 CAR T cell group significantly improved overall survival compared with other treatment groups (Figures 3C and 3G). In mice bearing SH-SY5Y, the combination ^177^Lu-NM600 and B7-H3 CAR T cell group showed significantly lower tumor burden than all other treatment groups (Figure 3H-I). Recipients of B7-H3 CAR T cells + ^177^Lu-NM600 RPT did not demonstrate visible signs of toxicity including skin rash or weight loss in either tumor model (Supplemental Figure S7A and B). Overall, these data indicate that RPT enhances the efficacy of CAR T cell therapy in two separate metastatic neuroblastoma xenograft models.

### Low-dose RPT is not detrimental to CAR T cells in the TME in vivo

As CAR T cell efficacy is related to its ability to traffic to tumor sites, we next examined CAR T cell trafficking in the context of low-dose RPT *in vivo*. In brief, NRG mice were intravenously injected with 1 x 10^6^ CHLA-20 or SH-SY5Y cells, then received either no treatment, B7-H3 CAR T cells, or B7-H3 CAR T cells + ^177^Lu-NM600 (Figure 4A). Fluorescently-labeled CAR T cell trafficking was observed 3 hours post-CAR T injection and every 24 hours for the first 3 days by fluorescence imaging (FLI). In both tumor models, the FLI revealed that CAR T cells are found initially and then remain primarily in the liver (Figure 4B and C), which corresponds to the location where the metastases are concentrated (as shown by the BLI imaging). In both tumor models, no difference was observed between CAR T cell trafficking in non-irradiated or irradiated conditions (Figure 4D and E; Supplemental Figure S8A and B).

**Figure 4.**
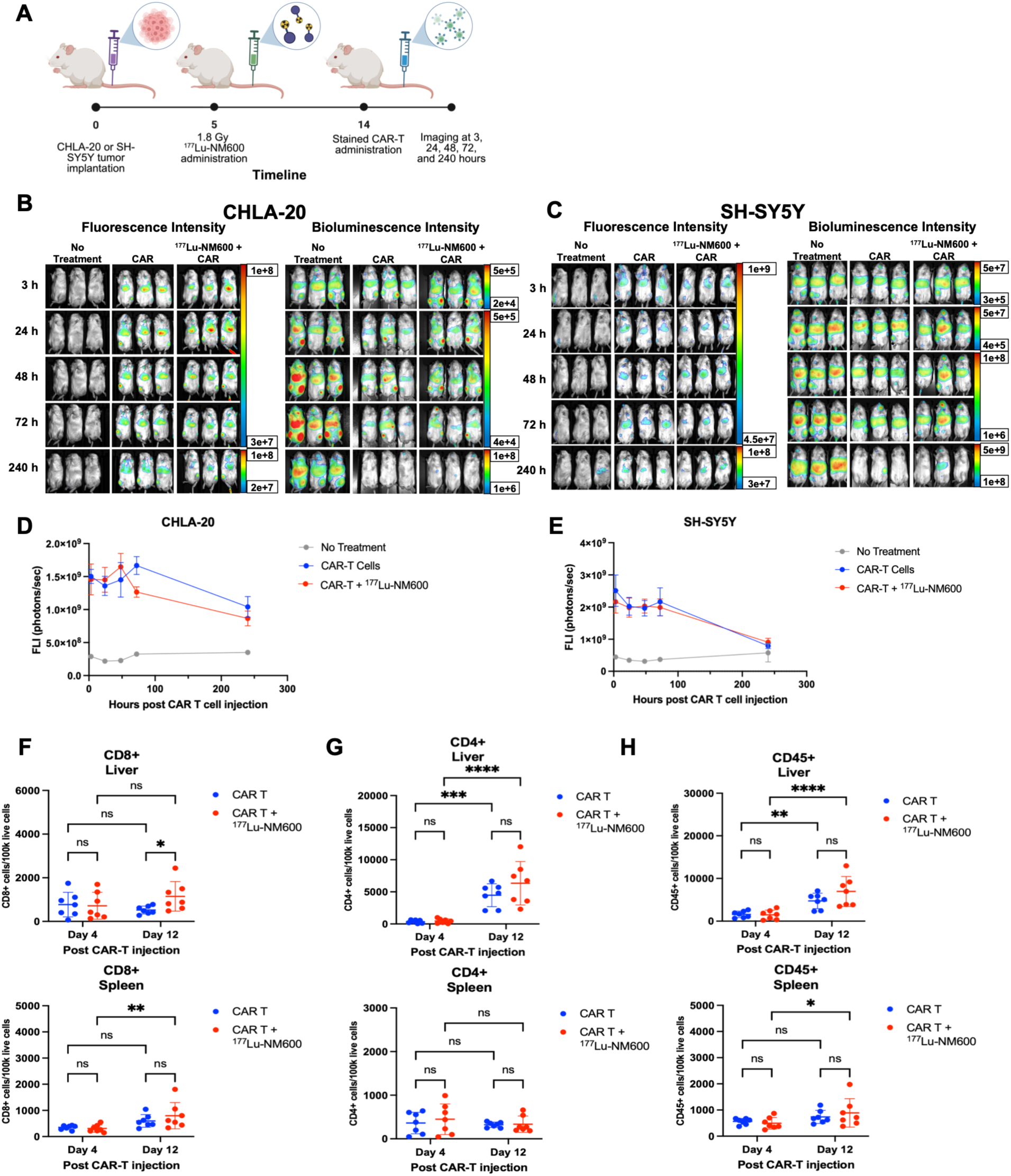
Low-dose RPT potentiates αB7-H3 CAR T cell residency despite minimal effects in trafficking *in vivo*. **A.** Schema of experimental design. **B-C.** Three representative Lago images of **(B)** CHLA-20 and **(C)** SH-Sy5Y bearing mice. CAR T cell fluorescence intensity (FLI) is displayed on the left and tumor burden bioluminescent intensity (BLI) on the right. Both measurements are displayed in photons/s. **D-E.** Line graph representing quantification of CAR T cell infiltration into **(D)** CHLA-20 and **(E)** SH-SY5Y tumor-bearing liver. Area under the curve of the FLI was evaluated for each group (CHLA-20: n = 3, no treatment, n = 4 CAR-T, and n = 5 CAR-T + ^177^Lu-NM600; SH-SY5Y: n = 5, all groups). Dunnett’s T3 multiple comparisons test was used to determine significance on the AUC. **F-H.** Flow cytometry to assess CAR T cell infiltration in a separate experiment using SH-SY5Y only as in (A), but spleen and liver were collected on day 4 and day 12 post-CAR T injection. **(F)** CD8+ in CD45+ CAR T **(G)** CD4+ cells in CD45+ CAR T cells and **(H)** human CD45+ CAR T cells were quantified. N = 7 mice per group were analyzed by a 2-way ANOVA. Significance is indicated as: ns (not significant), *\* (p < 0.05), ** (p < 0.01), *** (p < 0.001). ****(p < 0.0001)*.

To further characterize infiltrating CAR T cells in the absence or presence of RPT in a SH-SY5Y metastatic neuroblastoma xenograft model, we quantified human CD8^+^, CD4^+^, and CD45^+^ CAR T cell infiltration into the tumors and spleens of non-irradiated and irradiated mice by flow cytometry 4 and 12 days after CAR T administration (Figure 4F-H, Supplemental Figure S8C). In the liver, at day 12 post-B7-H3 CAR T cell infusion, an increase in CD8^+^ CAR T cells was detected after combination CAR T cells + ^177^Lu-NM600 compared to CAR T cells alone (Figure 4F). In the spleen, there was a significant increase in the number of CD8^+^ CAR T cells between day 4 and day 12 in the CAR T cell + ^177^Lu-NM600 group but not the CAR T cell treatment group (Figure 4F).

No significant differences were observed in the number of CD4^+^ CAR T cells between the CAR T and the combination group in either the liver or spleen. However, there was a significant increase in the number of CD4^+^ CAR T cells in the liver between day 4 and day 12 in both treatment groups (Figure 4G). These findings suggest that, regardless of radiation exposure, CD4-expressing CAR T cells may colonize the liver following CAR T therapy in metastatic neuroblastoma xenograft models.

We quantified human CD45^+^ cells (CAR T cells) by both flow cytometry (Figure 4H) and IHC (Supplemental Figure S8D-F) in the livers and spleens of mice bearing SH-SY5Y metastatic neuroblastoma. We detected fewer CD45^+^ cells in liver tissues in the absence of SH-SY5Y metastasis (Supplemental Figure S8D and E). No significant difference was detected between treatment groups in CD45^+^ cells in liver metastases, liver, or spleen. CD45^+^ cells were also detected in the spleens of mice from both treatment groups, with residence in splenic follicles.

Importantly, these data reveal that while B7-H3 CAR T cell trafficking is not increased by RPT, there are also no detrimental effects of the residual radiation dose delivered by RPT in the tumor, suggesting that CAR T cells can remain and potentially proliferate in a low-dose irradiated environment while retaining their efficacy.

### Low-dose RPT augments tumor cell expression of death receptors and immune susceptibility markers *in vitro*

The increased *in vivo* efficacy of RPT + CAR T cell treatment (Figure 3) despite no increase in CD45^+^ cell trafficking by RPT (Figure 4) suggests that the efficacy of this regimen is not defined by the number of CAR T cells present in the environment but potentially by the radiation-induced functionality of infiltrating CAR T cells or alterations to target cells. We therefore investigated whether low-dose radiation augmented the expression of death receptors and immunomodulatory markers on target cells, potentially affecting susceptibility to CAR T cell effector functions.

We quantified the effect of low-dose RPT on tumor cell expression of the death receptors Fas (CD95) (40) and TRAIL-R1 (CD261) (41), as well as immunosuppressive proteins PD-L1, HLA-DR, and Galectin-9. We also analyzed the poliovirus receptor (CD155), which serves a dual role in T cell regulation. T cells preferentially bind CD155 on tumor cells expressing co-inhibitory receptors TIGIT or CD96, but can also bind with an activating receptor, DNAM-1 (42). CHLA-20 and SH-SY5Y cells were incubated for 3 days with free ^177^Lu to achieve an estimated absorbed tumor cell dose of 2 Gy, followed by flow cytometry analysis (Figure 5A-F, Supplemental Figure 9). In CHLA-20 cells, the percentage of cells expressing Fas increased significantly after treatment with 2 Gy of ^177^Lu (Figure 5A). A small but significant decrease in the expression of PD-L1 was also observed (Figure 5B). In SH-SY5Y cells, Fas and TRAIL-R1 expression increased following treatment with 2 Gy ^177^Lu (Figure 5C) along with a modest increase in HLA-DR in SH-SY5Y cells (Figure 5D).

**Figure 5.**
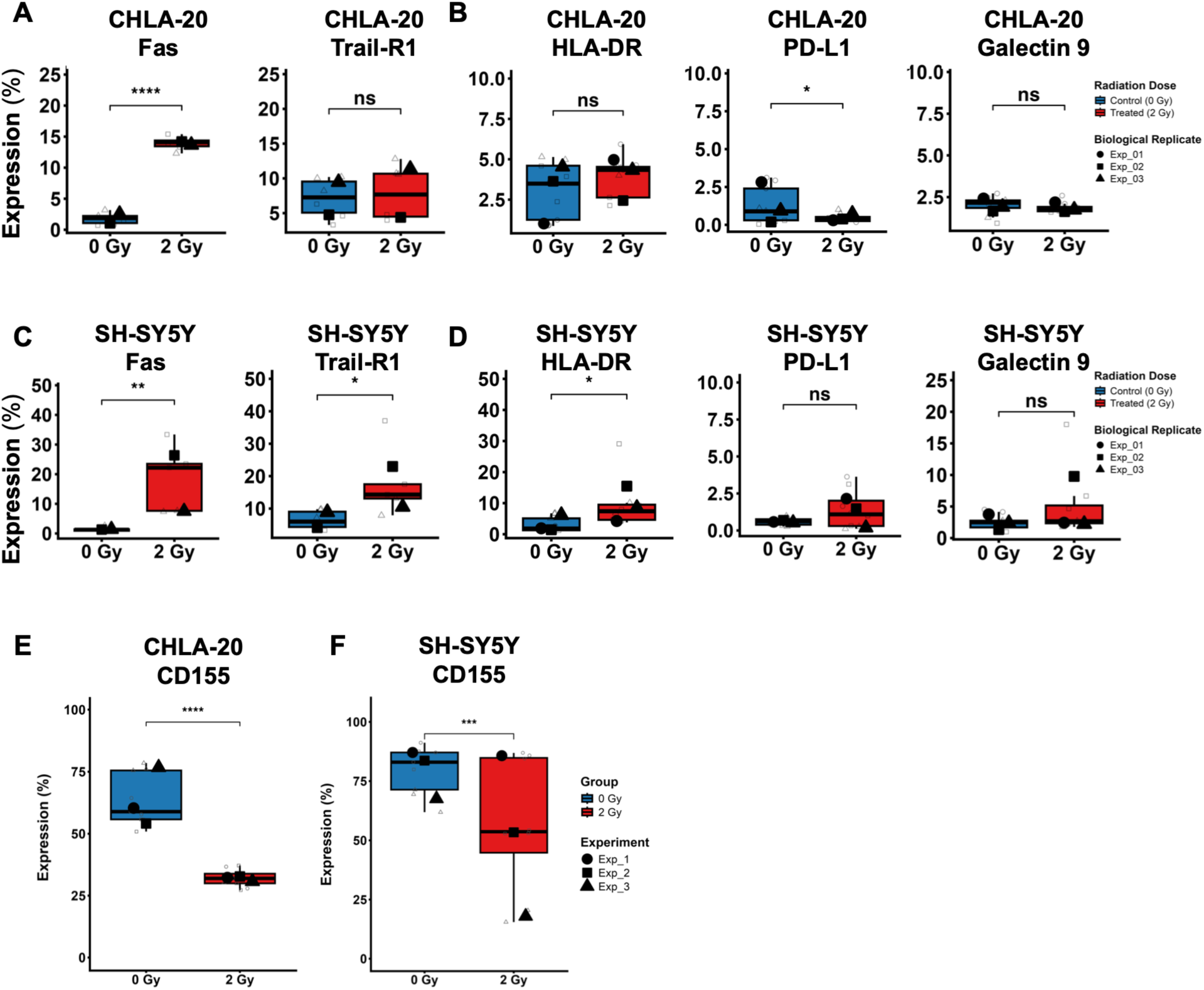
RPT induces pro-apoptotic but not immune susceptibility markers *in vitro.* CHLA-20 or SH-SY5Y cells were incubated with free ^177^Lu to reach a dose of 2 Gy after 3 days. On day 3, tumor cells were harvested and assessed by flow cytometry for **(A, C)** Fas and Trail-R1; **(B, D)** HLA-DR, PD-L1, Galectin 9; and **(E, F)** CD155 in the indicated cell line. Data are presented as a box and whisker plot. The biological experiment averages are displayed as a superplot, where large symbols represent the means of independent biological experiments (N = 3) and small symbols represent individual technical replicates (n = 3). Statistical significance was determined using a linear mixed model with radiation dose as a fixed factor and the biological experiments as a random effect to account for the nested technical replicates. Significance is indicated as: ns (not significant), *\* (p < 0.05), ** (p < 0.01), *** (p < 0.001). ****(p < 0.0001)*.

Following the irradiation of CHLA-20 and SH-SY5Y cells with 2 Gy of ^177^Lu, the expression of CD155, a co-inhibitory marker, was significantly decreased (Figure 5E and 5F). Collectively, these data suggest that low-dose ^177^Lu can alter phenotypic susceptibility of tumor cells that may enhance CAR T cell killing by selectively upregulating death receptor ligands such as Fas while downregulating inhibitor checkpoint ligands including CD155.

### Low-dose RPT-treated tumor cells promote functional resilience and sustained cytokine production in B7-H3 CAR T cells

Given that we did not consistently detect upregulation of immunosuppressive markers on neuroblastoma tumor cells following low-dose ^177^Lu, we next examined whether checkpoint markers (PD-1, Tim3, Lag3, TIGIT) on CAR T cells were altered after interaction with irradiated tumor cells. Compared to activated T cells, exhausted T cells produce little or no IL-2 and reduced TNFα, IFNγ, granzyme B, and perforin, which are critical to T cell proliferation and direct cytotoxic capability (43). To assess this, non-irradiated or irradiated CHLA-20 and SH-SY5Y cells were co-cultured with B7-H3 CAR T cells for 24 hours. Cytokines were quantified from the supernatant by ELISA, and PD-1, Tim3, Lag3, and TIGIT were quantified on CAR T cells by flow cytometry.

When co-cultured with CHLA-20 cells, no significant differences were observed in the secretion of the critical cytotoxic cytokines IFNγ, Granzyme B, and perforin (Figure 6A and 6B) while FasL, Granzyme A, IL-2, and TNFα were significantly reduced in co-cultures with irradiated tumor cells compared to non-irradiated tumor cells (Figure 6A, Supplemental Figure S10A and B). Irradiation significantly decreased the immunosuppressive cytokines IL-10 and IL-4 (Supplemental Figure 10C and D). This suggests that CAR T cells targeting CHLA-20 may have retained their effector machinery. Moreover, the attenuation of IL-4 and IL-10 suggests that radiation may sensitize CHLA-20 for a more sustained, less inhibited CAR T cell response.

**Figure 6.**
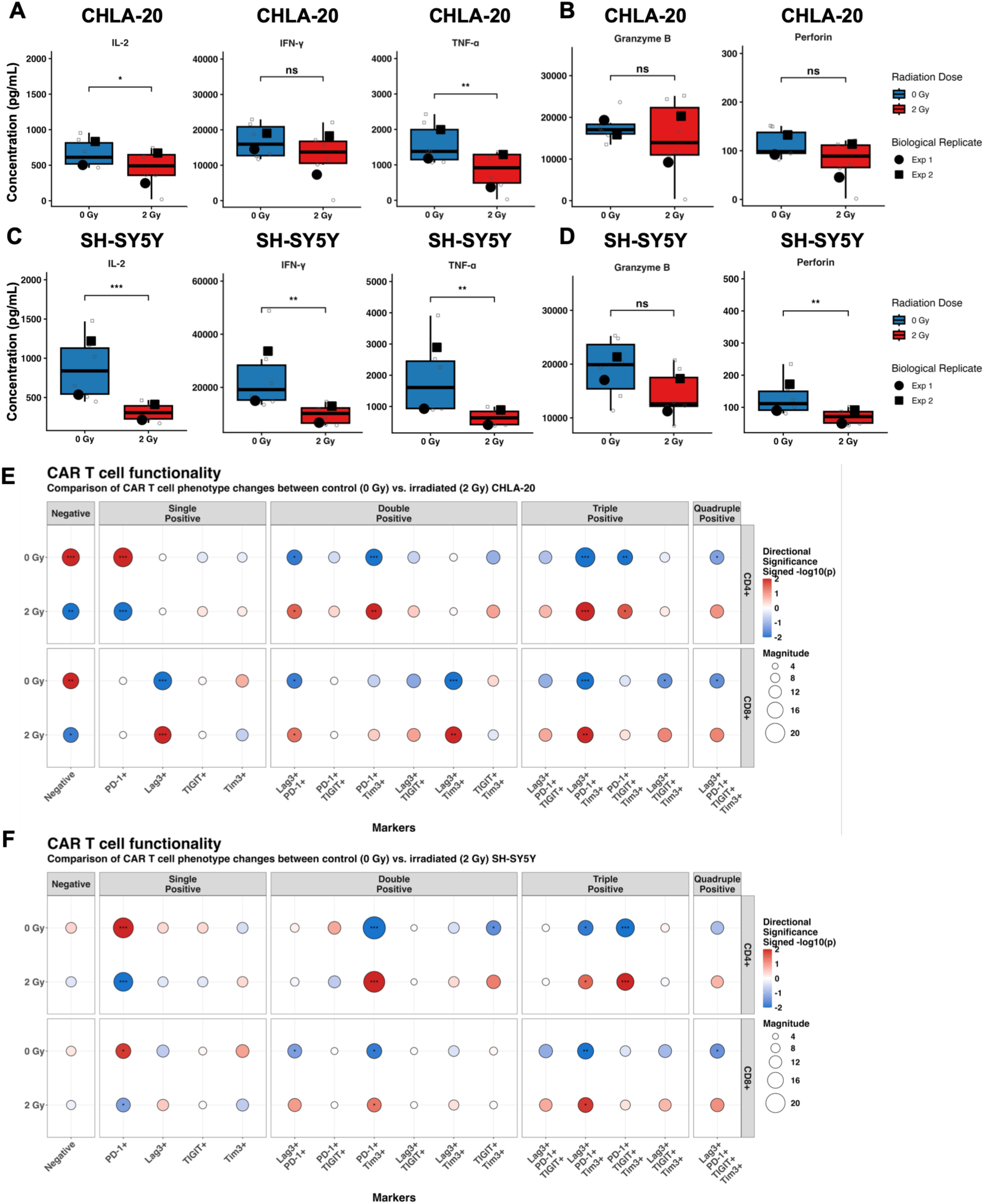
αB7-H3 CAR T cell functionality is not hindered by RPT. CHLA-20 or SH-SY5Y were plated and irradiated for 3 days with ^177^Lu to reach 2 Gy. On day 3, αB7-H3 CAR T cells were added in a 1:1 ratio and co-cultured for 24 hours. **A-D.** Concentration of the indicated cytokine in supernatant after 24 hours of co-culture with **(A-B)** CHLA-20 or **(C-D)** SH-SY5Y. Data are presented as a box and whisker plot. Large symbols represent the means of independent biological experiments (N = 2) and small symbols represent individual technical replicates (n = 3). Statistical significance was determined using a linear mixed model with radiation dose as a fixed factor and the biological experiments as a random effect to account for the nested technical replicates. Statistical significance was determined using Satterthwaite’s approximation for degrees of freedom. **E-F.** Pearson residuals analysis of the multi-checkpoint status of CD4+ (top) and CD8+ (bottom) CAR T cells (Donor 2) when co-cultured with **(E)** CHLA-20 or **(F)** SH-SY5Y for 24 hours. The size of the circle represents the magnitude of the Pearson residual, indicating the degree of deviation between the expected Vs. observed counts for each phenotype. Circle color denotes the directional significance (signed –log10(p)), where red represents enrichment and blue indicates depletion of a specific phenotype relative to the total population. Significance is indicated as: ns (not significant), *\* (p < 0.05), ** (p < 0.01), *** (p < 0.001). ****(p < 0.0001)*.

CAR T cell responses to irradiated SH-SY5Y were characterized by a significant and global suppression of the effector secretome at 24 hours (Figure 6C). This was associated with reductions in other cytotoxic molecules, such as perforin (Figure 6D), FasL (Supplemental Figure S10E), and Granzyme A (Supplemental Figure S10F). A significant reduction was also noted in IL-10 and IL4 (Supplemental Figure 10G and H).

In addition to cytokine secretion, we quantified the multi-checkpoint markers PD-1, Lag3, Tim3, and TIGIT in both CD8^+^ and CD4^+^ T cells after co-culture with ^177^Lu-treated or untreated CHLA-20 and SH-SY5Y cells using flow cytometry (Figure 6E and F, Supplemental Figure S11A and B). A Pearson residual bubble plot was used to quantify changes in T cells expressing single-positive (one marker), double-positive (two markers), triple-positive (three markers), or quadruple-positive (4 markers) exhaustion markers, also denoted as compartments. CD4+ and CD8+ populations from two donors were analyzed for the direction (circle color) and magnitude (circle size) of change in multi-checkpoint markers.

In the CD4^+^ CAR T population, the single positive PD-1 population was significantly decreased following co-culture with irradiated CHLA-20 cells (Figure 6E). Conversely, CAR T cells targeting non-irradiated CHLA-20 cells were significantly enriched for PD-1^+^ cells, while having minimal effects on the other markers within the single positive compartment. Additionally, significant enrichment in double-positive PD-1^+^Tim3^+^ and Lag3^+^PD-1^+^ CD4^+^ T cells was observed when co-cultured with irradiated CHLA-20 cells (Figure 6E). We also observed significant enrichment of Lag3^+^PD-1^+^Tim3^+^ and PD-1^+^TIGIT^+^Tim3^+^ populations within the triple-positive compartment.

Parallel analysis of the CD8^+^ CAR T cells revealed a transition toward a cytotoxic phenotype (Figure 6E). Following the co-culture of irradiated CHLA-20 cells (2 Gy delivered by ^177^Lu), there was a significant decrease of CD8^+^ CAR T cells expressing no exhaustion markers in the negative compartment. By comparison, the negative population in the CAR T cells co-cultured with non-irradiated CHLA-20 was increased. Furthermore, in the double positive compartment, CAR T cells targeting irradiated CHLA-20 shifted towards the Lag3^+^PD-1^+^ and Lag3^+^Tim3^+^ state. As seen in the CD4^+^ population, there was enrichment in the triple positive Lag3^+^PD-1^+^Tim3^+^ phenotype after co-culture with irradiated CHLA-20. Importantly, this transition was not detrimental to CAR T cell function as they maintained stable secretion of lytic cytokines when co-cultured with irradiated CHLA-20 cells (Figure 6A and B).

CD4^+^ CAR T cells responding to irradiated SH-SY5Y cells exhibited a significant decrease in PD-1^+^ single positive cells, while expanding on several multi-checkpoint phenotypes. We observed a significant enrichment of a PD-1^+^Tim3^+^ co-expressing CAR T cells. Additionally, radiation promoted triple positive Lag3^+^PD-1^+^Tim3^+^ and PD-1^+^TIGIT^+^Tim3^+^ phenotypes in CD4^+^, suggesting a maturation into a multi-checkpoint phenotype (Figure 6F).

## Discussion

Our previous study provided the initial evidence for combining CAR T cell therapy with low-dose RPT in a localized tumor setting (17). In this study, we aimed to evaluate if this combination would enhance anti-tumor responses in models of metastatic disease, given that CAR T cell therapy and RPT would most likely provide the highest clinical benefit in metastatic settings. We found combination B7-H3 CAR T cell therapy with ^177^Lu-NM600 RPT increased overall survival in two different metastatic neuroblastoma xenograft models as compared to monotherapy treatments. Although significant, this survival benefit did not extend beyond 100 days, as observed in our previous studies of localized flank tumor models (17). Nonetheless, these findings recapitulate our previous findings that an RPT regimen can safely enhance CAR T cell function against solid tumors, facilitating rather than inhibiting CAR T cell activity (17). Based on our prior data, we expect that the dose of RPT and the timing of CAR T cell administration are critical for this cooperative therapeutic effect.

This study dissected the potential mechanisms underlying increased survival in metastatic neuroblastoma models when CAR T cells and ^177^Lu-NM600 are given in combination. Fas was significantly increased on both CHLA-20 and SH-SY5Y cells, suggesting a CAR-independent killing mechanism in the enhanced CAR T cell function that may be involved in increasing the efficacy of CAR T cell treatment when combined with low-dose radiation delivered by RPT.

Immunomodulatory markers that might inhibit CAR T cell activity, including Galectin-9, and PD-L1, were largely unchanged on neuroblastoma after radiation. It has been shown that CD155 preferentially binds to TIGIT on T cells (42). Our observations suggest that radiation could potentially prevent CAR T cell exhaustion by significantly reducing the surface expression of CD155. Additionally, low-dose RPT was not detrimental to CAR T cell viability *in vivo* using our administration schedule, as reflected in CAR T cells numbers in the liver or spleen. This suggests that, in addition to directly killing tumor cells, low-dose RPT preserves the cytotoxicity of CAR T cells, providing a cooperative mechanism by which the therapeutic efficacy of the combination is enhanced, improving survival.

CAR T cells targeting both neuroblastoma models showed decreased cytokine production, although it is unclear whether these decreases affect efficacy. Cytokines are critical to cytotoxicity and cell renewal but are also associated with toxicities, such as cytokine release syndrome (44), suggesting moderation of cytokine release cold provide therapeutic benefit with lower morbidity. We observed a modest but significant population of multi-checkpoint Lag3^+^PD-1^+^TIGIT^+^ CD8^+^ and CD4^+^ cells. It is well established that terminally exhausted T cells produce less cytokines than their non-exhausted counterparts (43), which may in part account for decreased cytokines. However, given that low-dose RPT did not induce inhibitory immunomodulatory markers in CHLA-20 and SH-SY5Y, it is possible that this upregulation on CAR T cells did not trigger their functional exhaustion upon engagement of the tumor cells. However, these findings do suggest that our proposed therapeutic combination with CAR T cells and RPT in metastatic neuroblastoma may benefit from the addition of immune checkpoint inhibitors.

There are several limitations of our study. We used immunodeficient mice for the successful engraftment of human tumor cells. Unlike fully immune-competent models, immunodeficient mice lack certain mature endogenous lymphocyte populations, precluding assessment of other immune cell interactions. Initial work from Navarre et al in immunocompetent mice treated with CAR T cells and external beam radiotherapy suggests that activation of antigen presenting cells promotes activation and longevity CAR T cells (45), which can be explored in future studies adding RPT to CAR T cell therapy. Additionally, the diffuse nature of the metastatic tumors in the liver limited the isolation of CAR T cells and tumor cells co-localized *in vivo*. While we were able to study these interactions *in vitro*, exposure of cells to ^177^Lu *in vitro* likely differed compared to those exposed *in vivo*.

While we investigated only neuroblastoma models, B7-H3 is expressed on the surface of a myriad of other tumor types (38). Additionally, NM600 has pan-tumor targeting capabilities (16, 18, 21). Given the ability of RPTs to deliver radiation systemically to metastatic tumors, this combination has the potential to be used as a pan-cancer treatment in other metastatic diseases. Future studies might include additional tumor types as well as orthotopic or spontaneous tumor models in immunocompetent mice to fully evaluate the immunomodulatory effects of combining CAR T cells and RPT.

This study utilized the beta-particle emitter ^177^Lu. While we did observe significant differences in probability of survival and tumor reduction *in vivo*, beta-particle emitters have a medium path length and low linear energy transfer (LET). Optimization of these characteristics of the selected radionuclide is possible, given that NM600 and other targeting agents can be chelated to various isotopes. Future studies may elucidate whether CAR T combinations in metastatic disease are more optimal with a higher LET but shorter pathlength isotope, such as ^225^Ac.

This study demonstrates in metastatic neuroblastoma models not only that CAR T cells can remain and potentially proliferate in a low-dose RPT-irradiated environment, but that low-dose RPT enhances the efficacy of CAR T cell therapy. Increased survival was exhibited in two xenograft models of metastatic neuroblastoma compared to either therapy alone. Although these findings have the potential to be extended to other solid tumor models, additional studies must investigate for each model the timing and sequence of administration of RPT and CAR T cells and perform dosimetry studies for RPT agents to determine the optimal doses of RPT needed. Furthermore, elucidation of mechanisms driving anti-tumor response will be essential to fully evaluate the clinical potential of RPT in combination with CAR T cell therapy.

## Transparency declaration

The lead author* affirms that this manuscript is an honest, accurate, and transparent account of the study being reported; that no important aspects of the study have been omitted; and that any discrepancies from the study as planned (and, if relevant, registered) have been explained.

*The manuscript’s guarantor.

## Availability of data and material

The datasets used and/or analyzed during the current study are available from the corresponding author on reasonable request.

## Authors’ contributions

AGS: Conceptualization, methodology, software, formal analysis, writing—original draft, visualization, investigation.

AO: investigation, Writing—reviewing and editing

AW: investigation, Writing—reviewing and editing

LS: investigation, Writing—reviewing and editing

AKE: investigation, Writing—reviewing and editing

IHA: investigation, Writing—reviewing and editing

MBI: investigation, Writing—reviewing and editing

OK: investigation, Writing—reviewing and editing

HCR: investigation, Writing—reviewing and editing

MB: investigation, Writing—reviewing and editing

JA: Resources, Writing—reviewing and editing

JCE: Formal Analysis, Writing—reviewing and editing

IO: Formal Analysis, Data Curation, Supervision, Writing—reviewing and editing

RMR: Resources, Writing—reviewing and editing

PMS: Conceptualization, Supervision, Writing—reviewing and editing

CMC: Conceptualization, Supervision, Writing—reviewing and editing

RH: Resources, Writing—reviewing and editing

BPB: Resources, Writing—reviewing and editing

JPW: Resources, Writing—reviewing and editing

ZSM: Conceptualization, Supervision, Writing—reviewing and editing, Funding Acquisition, Project Administration, Data Curation

QHS: Conceptualization, Supervision, Writing—reviewing and editing, Funding Acquisition, Project Administration, Data Curation

## Acknowledgments

We thank T.J. Berg for the editorial review of this manuscript. We would also like to thank all the other collaborators who provided advise and expertise throughout the process of this manuscript. We also thank Google Gemini for the assistance in R code generation. We also thank the outstanding veterinary staff in the WIMR vivarium for the care of mice throughout in vivo studies, the University of Wisconsin Carbone Cancer Center (UWCCC) Flow Cytometry core facility, UWCCC Translational Research Initiatives in Pathology core facility, and the UWCCC Small Animal Imaging and Radiotherapy core facility for their expertise.

