## Supplemental Data for "Low-dose radiopharmaceutical therapy enhances the efficacy of B7-H3 CAR T cells in murine metastatic neuroblastoma"

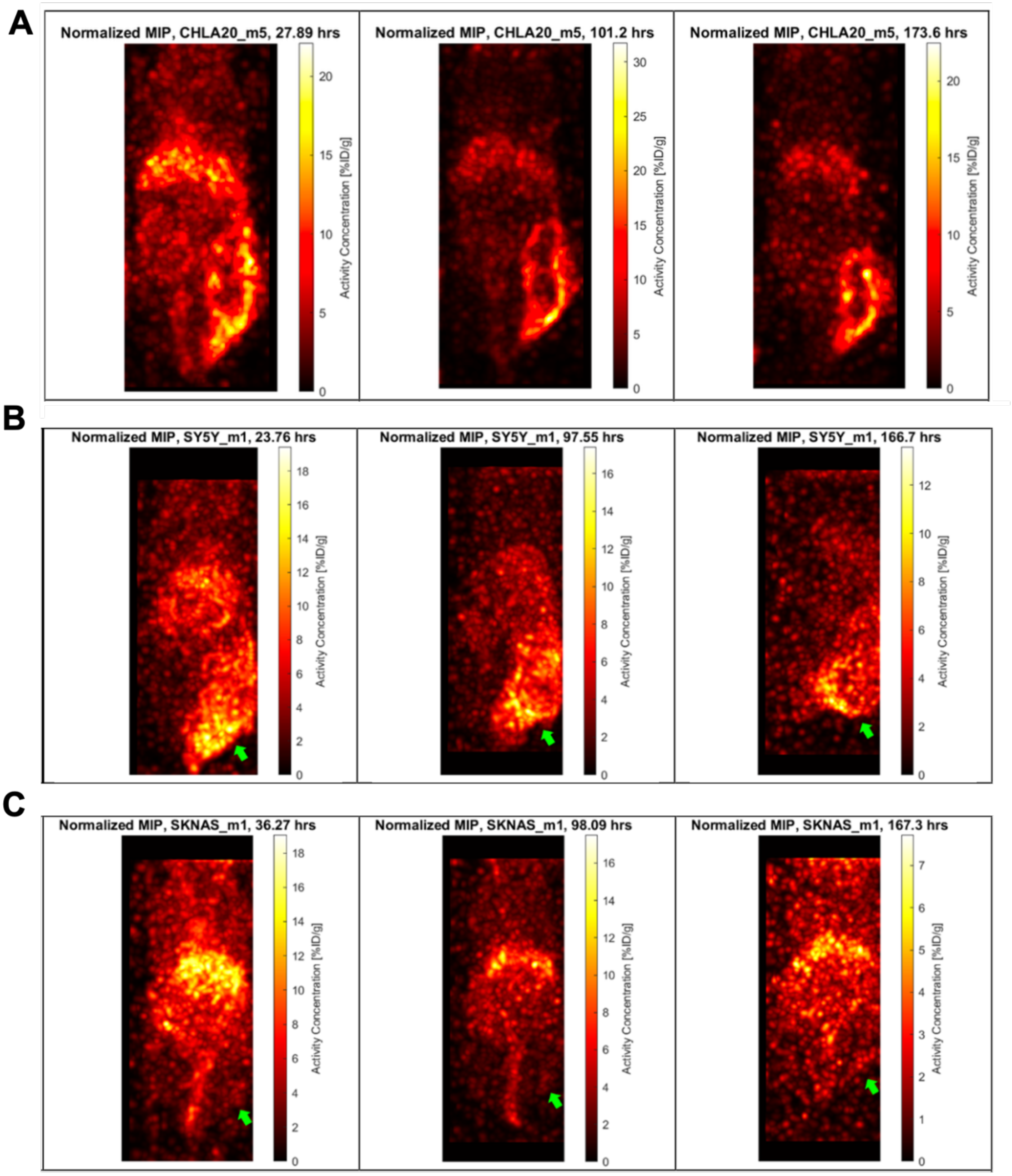


**Supplemental Figure S1. Maximum intensity projections (MIPs) in units of activity concentration (%ID/g) of three flank models of neuroblastoma. A.** Representative CHLA-20 tumor-bearing mouse over 3 time points showing uptake of $^{177}$Lu-NM600 to the tumor. **B.** Representative SH-SY5Y tumor-bearing mouse over 3 time points showing uptake of $^{177}$Lu-NM600 to the tumor. **C.** Representative SK-N-AS tumor-bearing mouse over 3 time points showing uptake of $^{177}$Lu-NM600 to the tumor.

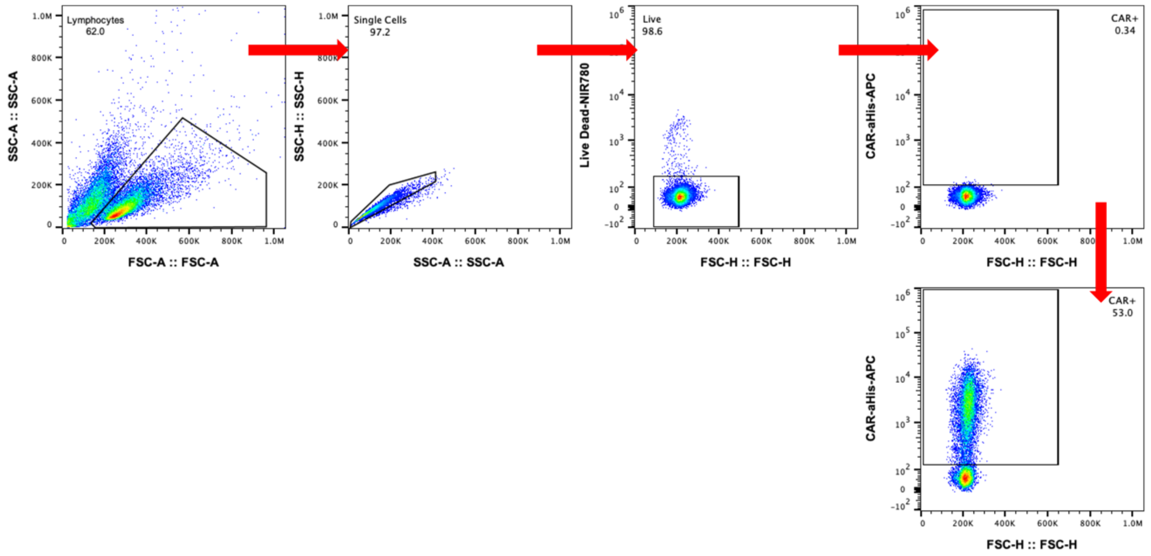


**Supplemental Figure S2. B7-H3 CAR T flow cytometry gating.** Gating strategy for CAR expression.

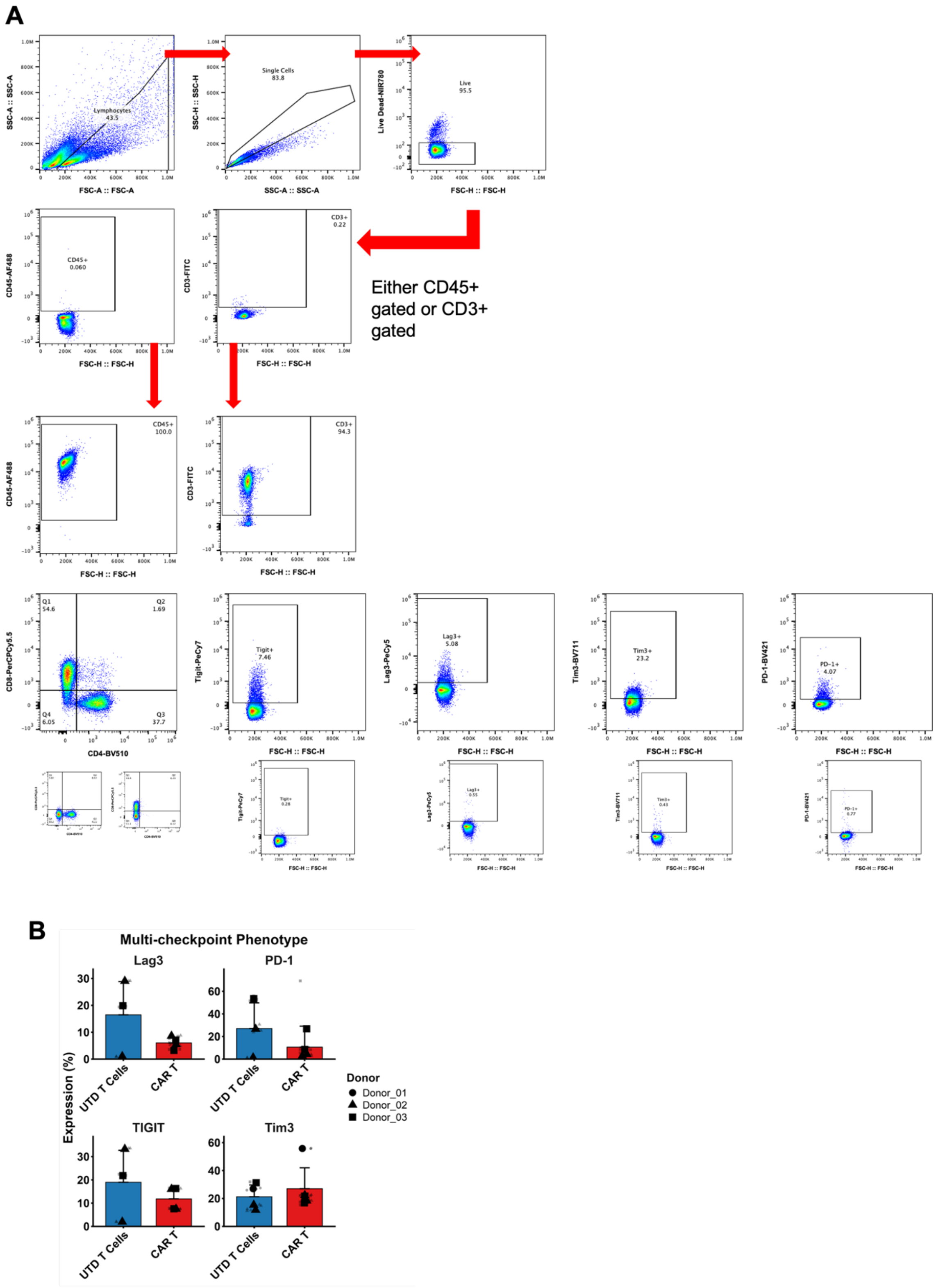


**Supplemental Figure S3. Multi-checkpoint markers on B7-H3 CAR T cells. A.** Gating strategy for CD8, CD4, and multi-checkpoint phenotype post thaw. **B.** Flow cytometry quantification of multi-checkpoint markers on CAR T cells and UTD T cells post-thaw. Circle represents Donor 1, square represents Donor 2, and triangle represents Donor 3. Data is represented as the grand mean ± SD of n = 3 technical replicates and either n = 2 (transduced) or n = 1 (untransduced) biological replicates.

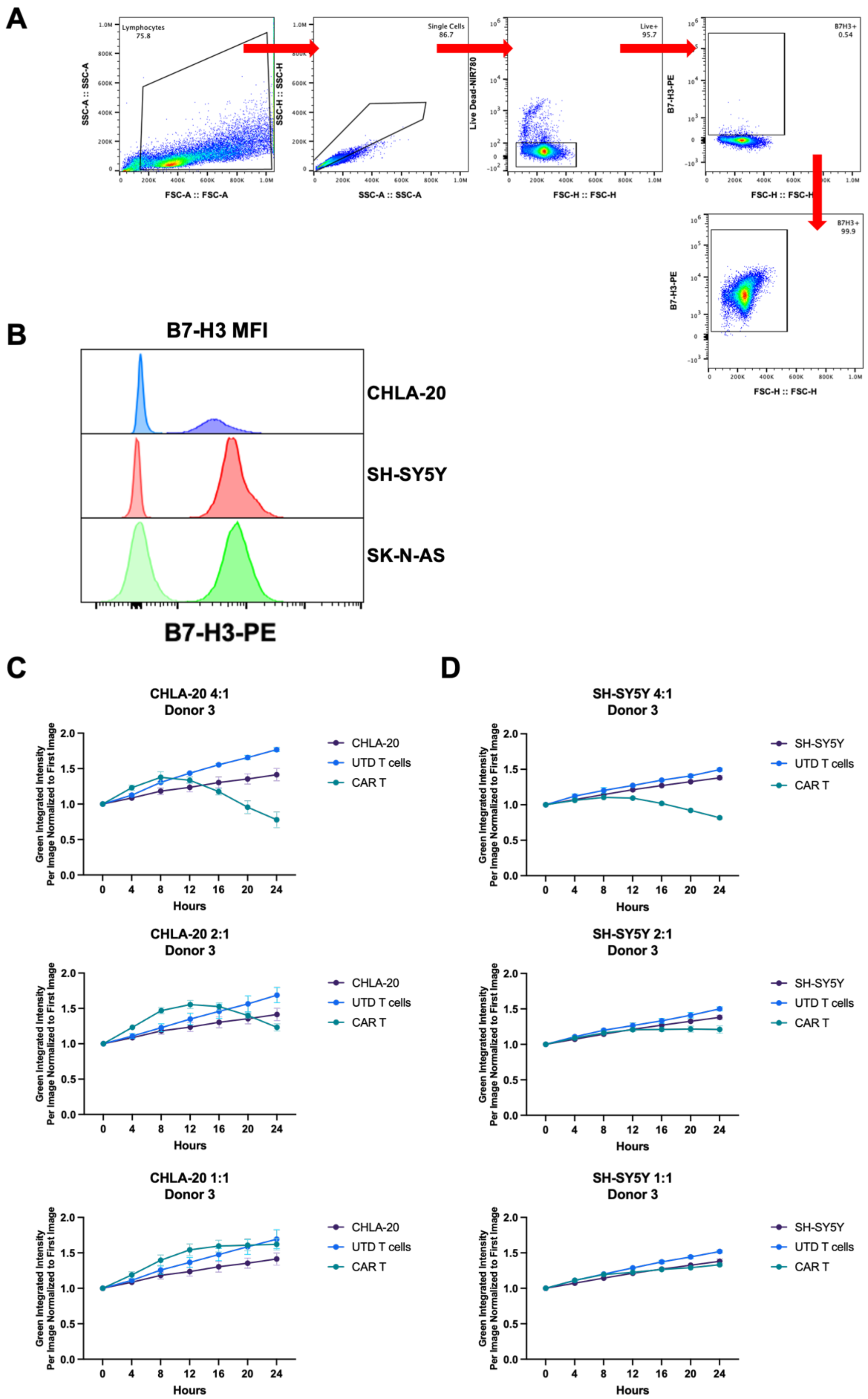


**Supplemental Figure S4. B7-H3 CAR T cells have cytotoxicity against multiple models of neuroblastoma. A.** Gating strategy for B7-H3$^+$ tumor lines CHLA-20, SH-SY5Y, and SK-N-AS. **B.** Representative histogram data of B7-H3 expression of CHLA-20, SH-SY5Y, and SK-N-AS. The first peak represents isotype staining. **C.** Cytotoxicity assay of Donor 3 CAR T cells targeting CHLA-20. Either UTD

T cells or CAR T cells were plated at 4:1, 2:1, or 1:1 effector-to-target (E:T) ratios against each of the tumor lines. Cells were co-cultured for 20+ hours, and a decrease in green fluorescence intensity was used to assess cytotoxic capabilities, normalized to the first image. Data represent one experiment of Donor 3 (N = 1, technical replicates n = 3). **D.** Cytotoxicity assay of Donor 3 CAR T cells targeting SH-SY5Y. Either UTD T cells or CAR T cells were plated at 4:1, 2:1, or 1:1 effector-to-target (E:T) ratios against each of the tumor lines. Cells were co-cultured for 20+ hours and a decrease in green fluorescence intensity was used to assess cytotoxic capabilities, normalized to the first image. Data represent one experiment of Donor 3 (N = 1, technical replicates n = 3).

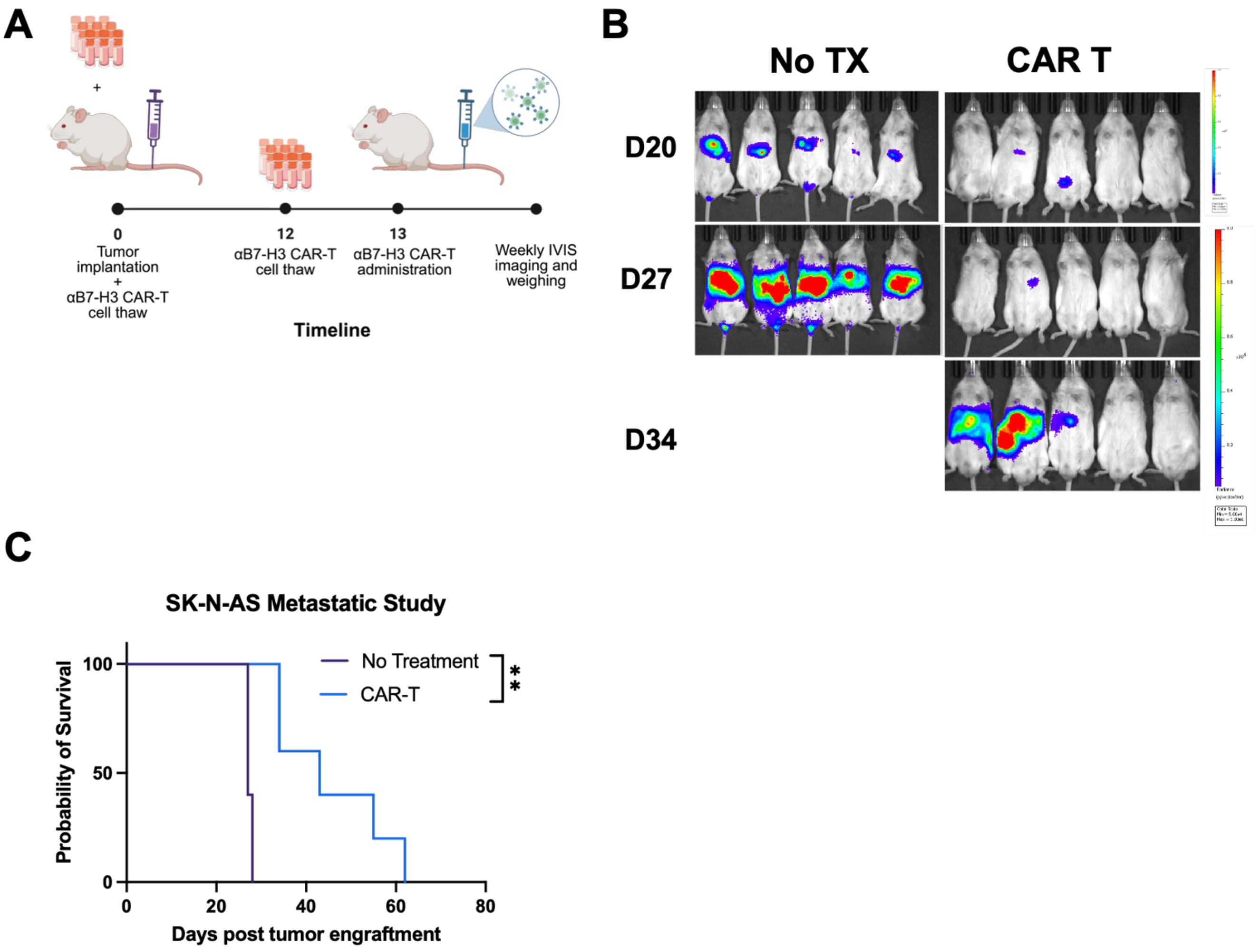


**Supplemental Figure S5. B7-H3 CAR-T efficacy in metastatic neuroblastoma model. A.** Schematic *in vivo* study of B7-H3 CAR-T cells targeting a metastatic model of SK-N-AS in NRG mice. Figure schema created in Biorender. 1 x $10^6$ SK-N-AS cells were tail vein injected into NRG mice. Mice were sorted into two groups: control (No Treatment: $n = 5$), and B7-H3 CAR T alone ($n = 5$; $4 \times 10^6$ CAR+ cells). The treatment effect was assessed by the probability of survival. Significance is indicated as ** *($p < 0.01$).*

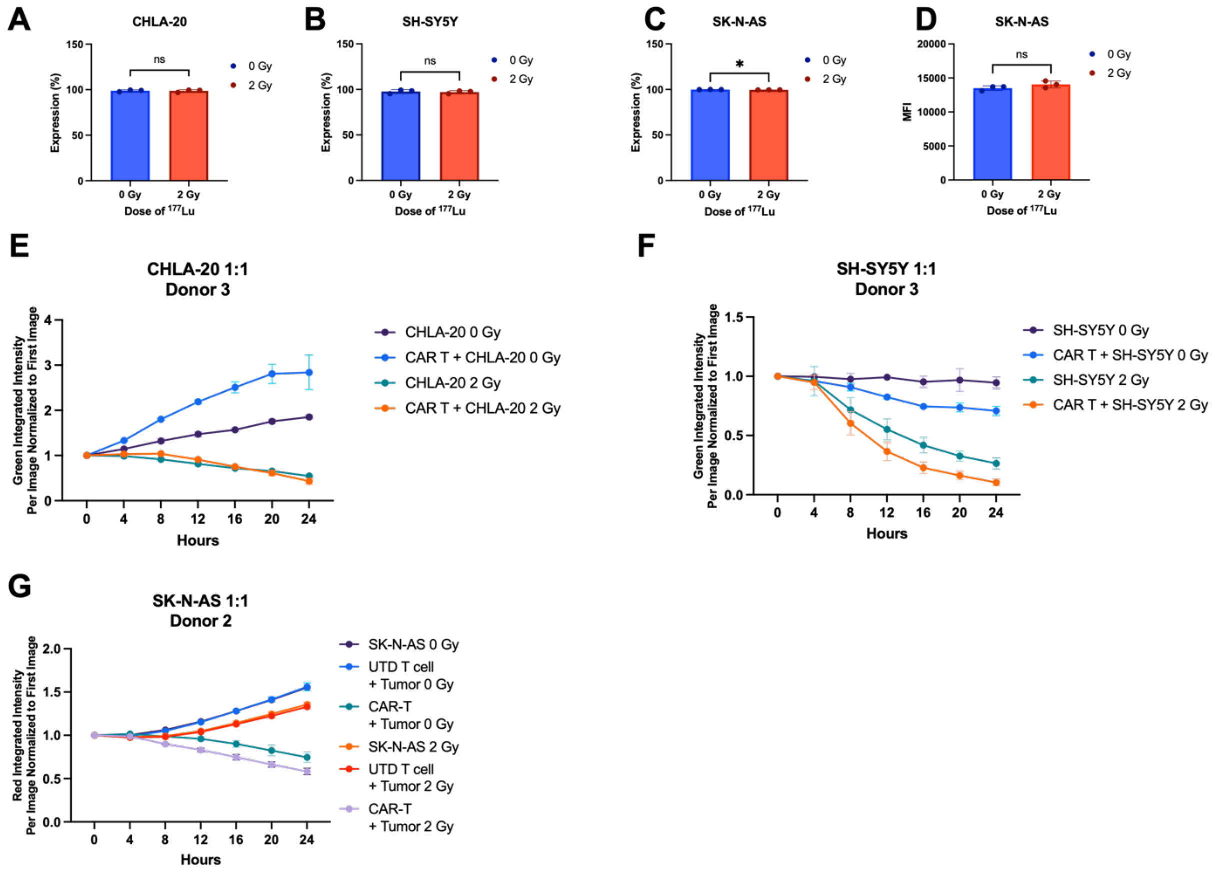


**Supplemental Figure S6. Low dose RPT enhances *in vitro* cytotoxicity. A-C.** Percent expression of B7-H3 on CHLA-20, SH-SY5Y, and SK-N-AS when incubated with 2 Gy $^{177}$Lu for 3 days. For CHLA-20 and SH-SY5Y, data are represented as the average ± SD of n = 3 independent experiments, each performed with n = 3 triplicates. **D.** B7-H3 MFI on SK-N-AS when incubated with 2 Gy $^{177}$Lu for 3 days. SK-N-AS data are represented as mean ± SD of n = 3 triplicates. **E.** Cytotoxicity assay of CHLA-20 targeted by Donor 3 B7-H3 CAR T and UTD T cells. Tumor cells were incubated with 1 Gy of $^{177}$Lu for 3 days. B7-H3 CAR T and UTD T cells were added in a 1:1 E:T ratio. **F.** Cytotoxicity assay of SH-SY5Y targeted by Donor 3 B7-H3 CAR-T and UTD T cells. Tumor cells were incubated with 2 Gy of $^{177}$Lu for 3 days. B7-H3 CAR-T and UTD T cells were added in a 1:1 E:T ratio. **G.** Cytotoxicity assay of SK-N-AS with Donor 2 B7-H3 CAR T and UTD T cells. Tumor cells were incubated with 2 Gy of $^{177}$Lu for 3 days. B7-H3 CAR T and UTD T cells were added in a 1:1 E:T ratio. Cytotoxicity assay data are presented as one donor's mean ± SD of n = 3 triplicates.

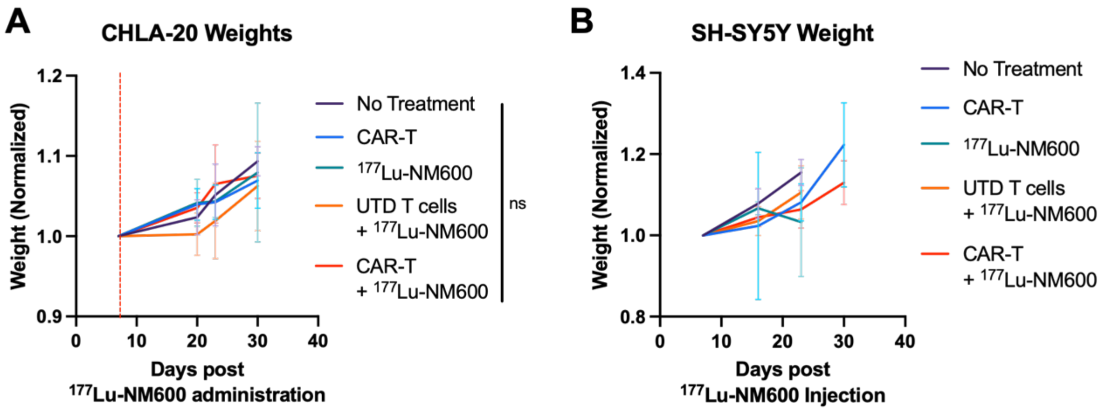


**Supplemental Figure S7. Mouse weights with treatment over time. A.** CHLA-20 weights in each group averaged over time. **B.** SH-SY5Y weights in each group averaged over time.

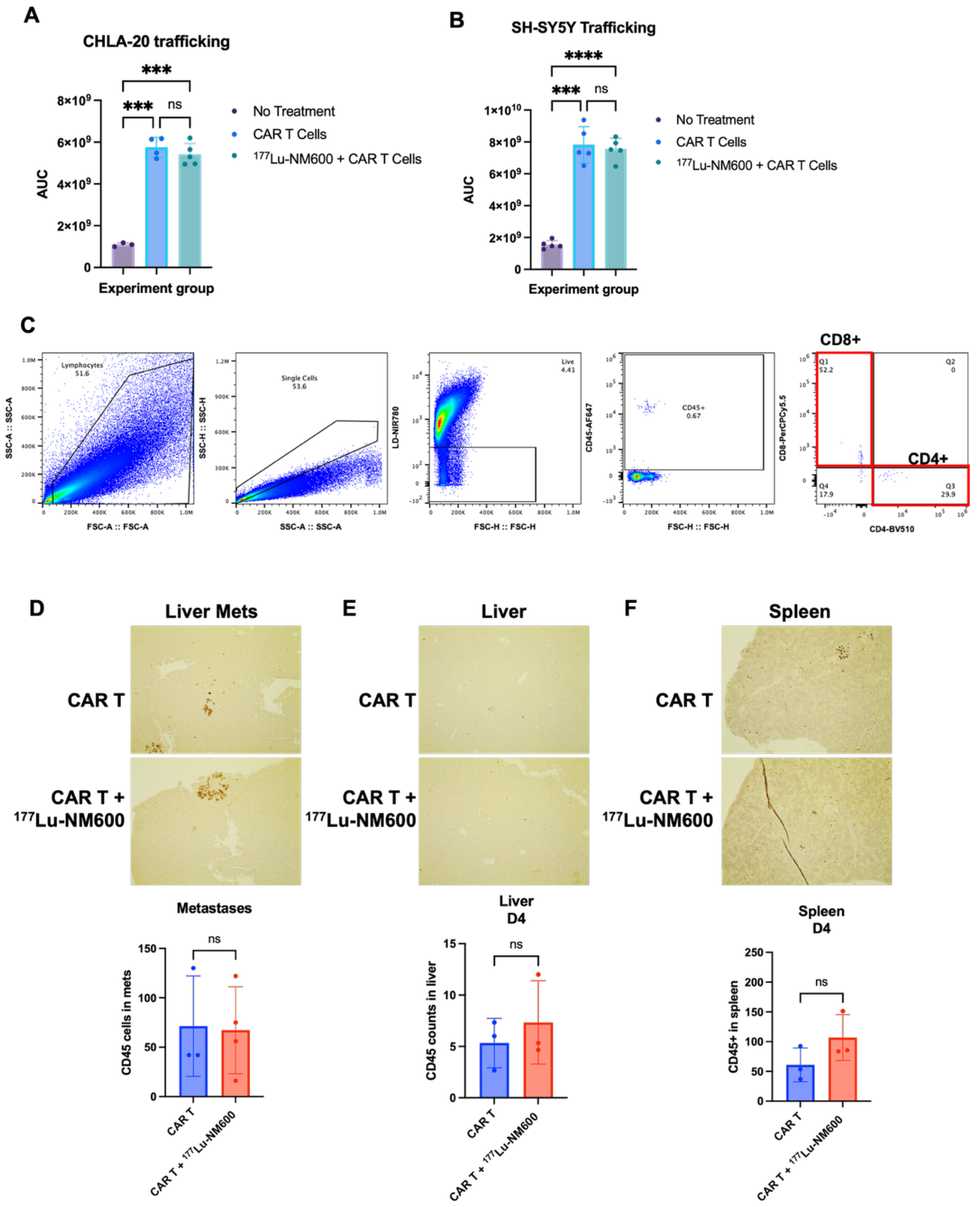


**Supplemental Figure S8. Low dose RPT does not inhibit B7-H3 CAR T cell infiltration. A.** Bar graph of the area under the curve (AUC) of the FLI of mouse imaging for CHLA-20. **B.** Bar graph of the area under the curve (AUC) of the FLI of mouse imaging for SH-SY5Y. Significance is indicated as: ns (not significant), * *($p < 0.05$), ** ($p < 0.01$), *** ($p < 0.001$). ****($p < 0.0001$).* **C.** Gating strategy for CD45, CD8, and CD4 expressing cells in the liver and the spleen. **D-F** Immunohistochemistry (IHC) images of human CD45 stained CAR T cells. **(D)** $CD45^+$ cells in liver metastasis of SH-SY5Y on day 4 after CAR T administration. No metastases were detected in the liver at day 12 after CAR T injection. Images were taken at 20X. CD45 expressing cells were counted in the metastases of each group. In CAR T group, n = 3 metastases were detected, while in the CAR T + $^{177}$Lu-NM600 n = 4 metastases were detected. **E.** IHC images of human CD45 CAR T cells in the liver on day 4 and day 12. Images were taken at 10X. CD45 cells were counted in 3 fields across the liver in 3 mice and averaged. **F.** IHC images of human CD45 CAR T cells in the spleen on day 4 and day 12. Images were taken at 10X. CD45 cells were counted in 3 fields across the spleen in 3 mice and averaged. A Welch's t-test was used to determine significance in IHC images.

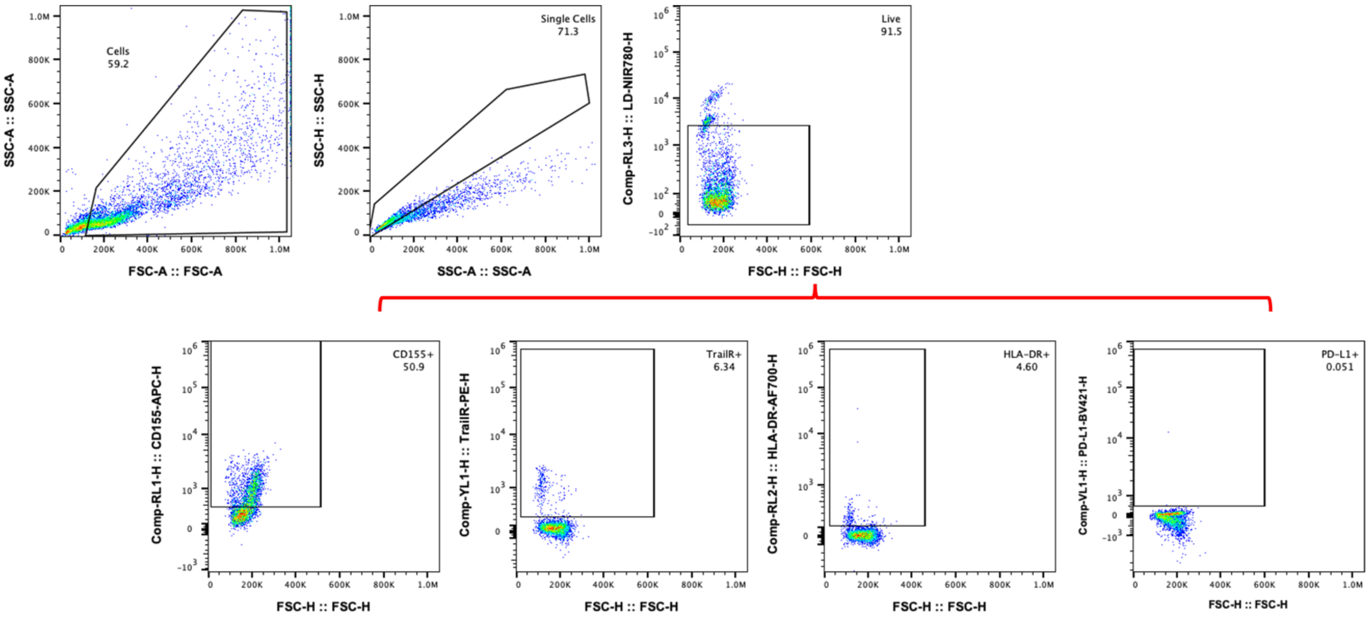


**Supplemental Figure S9. RPT induces pro-apoptotic but not immunomodulatory markers *in vitro.* A**. Gating strategy for Fas, Trail-R1, HLA-DR, PD-L1, Galectin 9, and CD155. Total cells were gated, followed by doublet cell exclusion. Live cells were then gated. Fluorescence minus one (FMO) was used to place the marker of interest gate within the live cells. Cells were derived from the CHLA-20 tumor model.

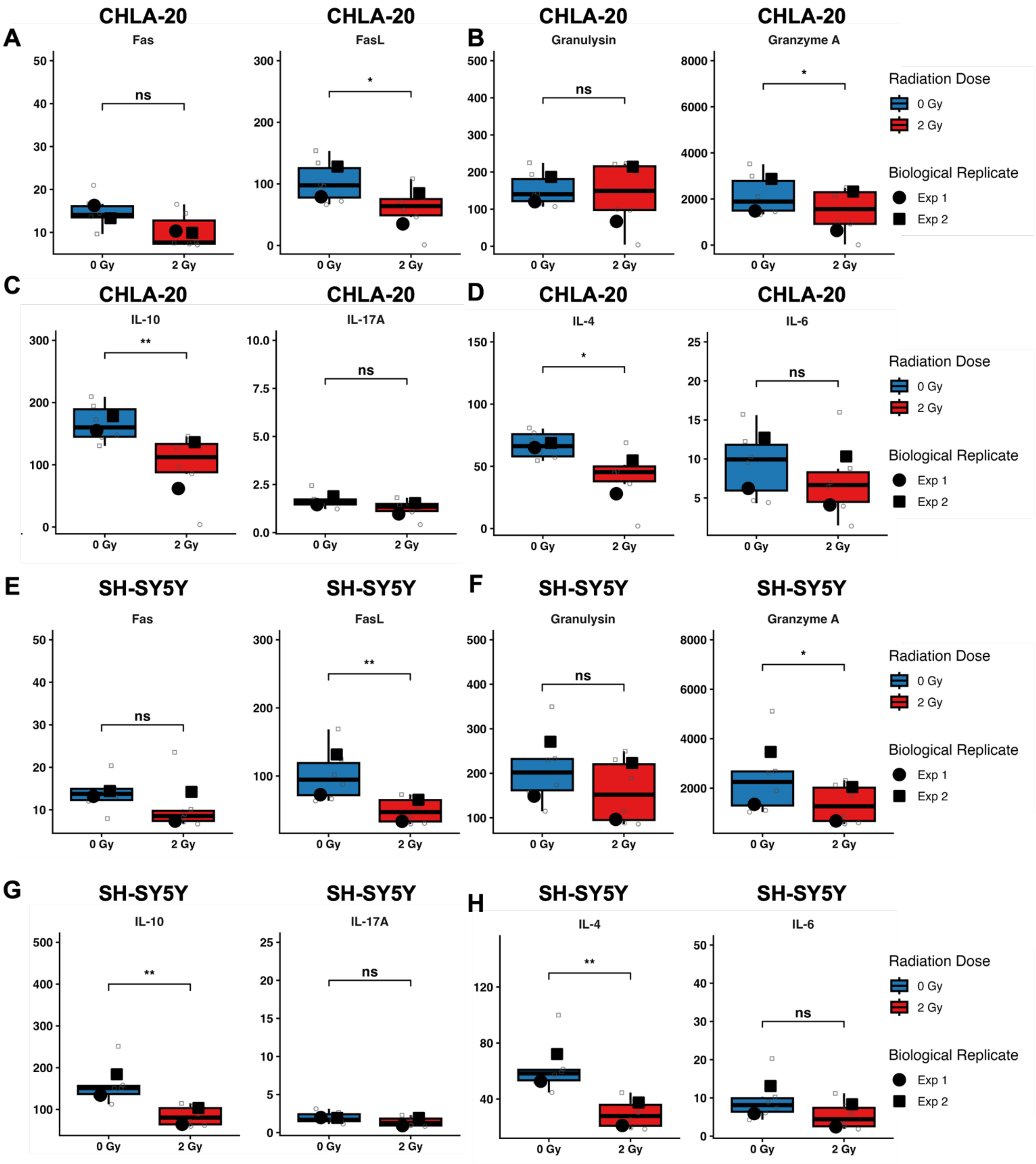


**Supplemental Figure S10. B7-H3 CAR T cytokine release.** **A-D.** Cytokine release of CAR T cells co-cultured with CHLA-20 or **E-H.** SH-SY5Y. Statistical significance was determined using a linear mixed model with radiation dose as a fixed factor and the biological experiments as a random effect to account for the nested structure of the data. Statistical significance was determined using Satterthwaite's approximation for degrees of freedom. Significance is indicated as: ns (not significant), * *($p < 0.05$)*, ** *($p < 0.01$)*, *** *($p < 0.001$)*. ****(*$p < 0.0001$*).

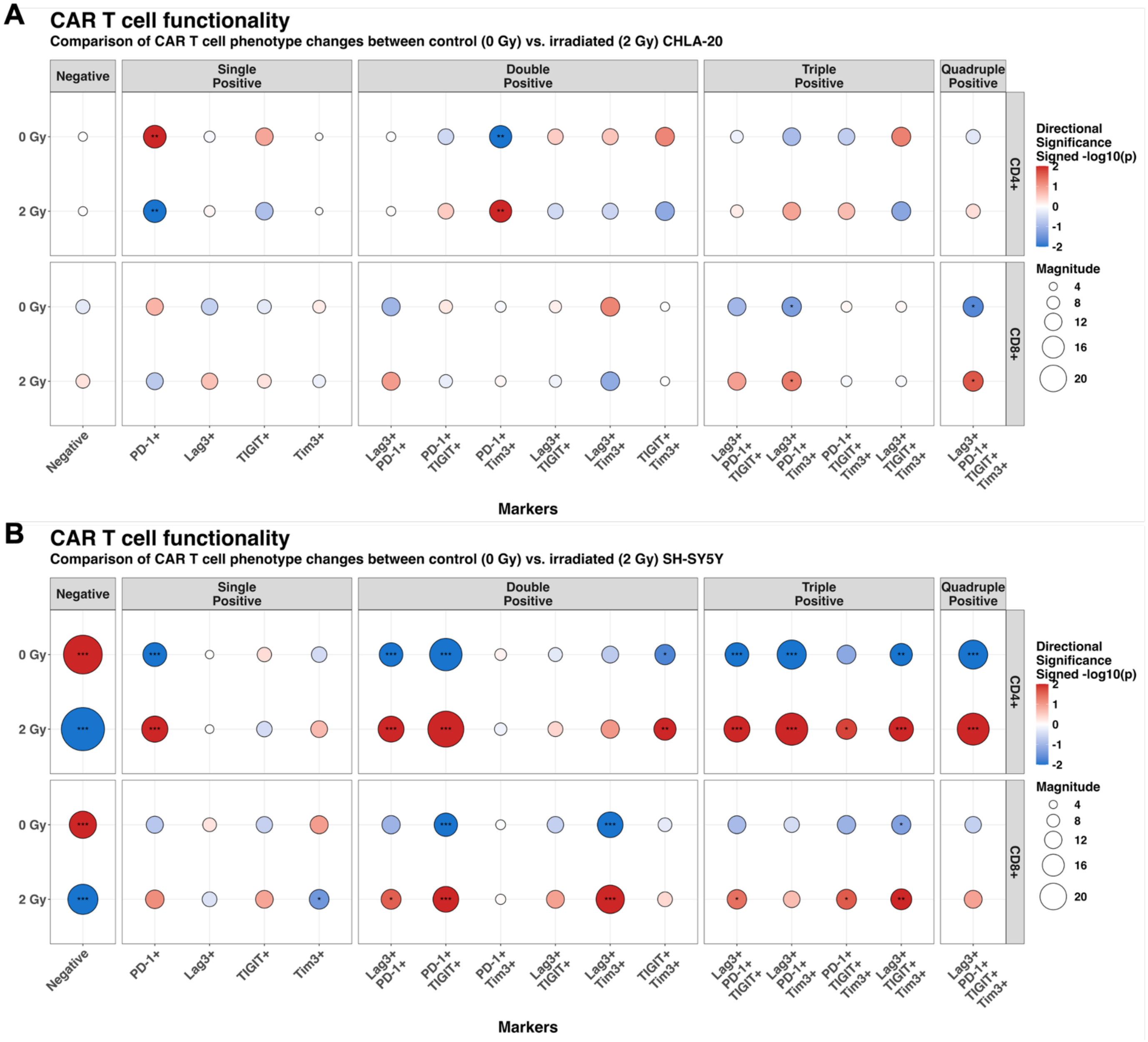


**Supplemental Figure S11. B7-H3 CAR T cytokine release. A-B.** Pearson residuals analysis of the multi-checkpoint status of CD4+ (top) and CD8+ (bottom) CAR T cells (Donor 3) when co-cultured with **(A)** CHLA-20 or **(B)** SH-SY5Y for 24 hours. The size of the circle represents the magnitude of the Pearson residual, indicating the degree of deviation between the expected vs observed counts for each phenotype. Circle color denotes the directional significance (signed –log10(p)), where red represents enrichment and blue indicates depletion of a specific phenotype relative to the total population. Significance is indicated as: ns (not significant), * *(p < 0.05), ** (p < 0.01), *** (p < 0.001). ****(p < 0.0001).*

**Supplemental Table S1:** *Antibodies used in the study.*

| Antibody | Clone | Catalog | Fluorophore | Vendor |
|---|---|---|---|---|
| CD8 | SK1 | 344710 | PerCPCy5.5 | Biolegend |
| Lag3 | 11C3C65 | 369346 | PeCy5 | Biolegend |
| Tigit | A15153G | 372714 | PeCy7 | Biolegend |
| PD-1 | EH122H7 | 329920 | BV421 | Biolegend |
| CD4 | SK3 | 344634 | BV510 | Biolegend |
| TIM3 | 7D3 | 565566 | BV711 | BD Biosciences |
| CD45 | HI30 | 304017 | AF488 | Biolegend |
| CD45 | 2D1 | 368538 | AF647 | Biolegend |
| Galectin 9 | 9M1-3 | 348916 | PeCy7 | Biolegend |
| PD-L1 | MiH3 | 374508 | BV421 | Biolegend |
| CD155 | SKIL4 | 337618 | APC | Biolegend |
| HLA-DR | LN3 | 327014 | AF700 | Biolegend |
| TrailDR | DJR1 | 307206 | PE | Biolegend |
| FAS | DX2 | 305644 | BV711 | Biolegend |
| anti-His Tag | J095G46 | 362605 | APC | Biolegend |
| anti-His Tag | J095G46 | 362603 | PE | Biolegend |
| Mouse IgG2a, κ Isotype Ctrl | MOPC-173 | 400212 | PE | Biolegend |
| Mouse IgG2a, κ Isotype Ctrl | MOPC-173 | 400220 | APC | Biolegend |
| Recombinant Human B7-H3 His-tag Protein | N/A | 1949-B3-050 | Primary | RD Systems |

**Supplemental Table S2:** ***R libraries used in the study.***

| R Libraries | Reference |
| --- | --- |
| library(CytoML) | Jiang, M. CytoML. Bioconductor https://doi.org/10.18129/B9.BIOC.CYTOML (2017). |
| library(readxl) | Wickham, H. & Bryan, J. Readxl: Read Excel Files. (2023). |
| library(lme4) | Bates, D., Mächler, M., Bolker, B. & Walker, S. Fitting Linear Mixed-Effects Models Using lme4. Journal of Statistical Software 67, 1–48 (2015) |
| library(lmerTest) | Kuznetsova, A., Brockhoff, P. B. & Christensen, R. H. B. lmerTest Package: Term Tests and Degrees of Freedom Methods for Linear Mixed-Effects Models. Journal of Statistical Software 82, 1–26 (2017). |
| library(emmeans) | Lenth, R. V. Emmeans: Estimated Marginal Means, Aka Least-Squares Means. (2024). |
| library(biostats) | Quirarte-Justo, S., Montano-Ruiz, A. C. & Torres-Arellano, J. M. Biostats: Biostatistics and Clinical Data Analysis. (2026). |
| library(ggplot2) | Wickham, H. Ggplot2: Elegant Graphics for Data Analysis. (Springer-Verlag New York, 2016). |
| library(ggsignif) | Ahlmann-Eltze, C. & Patil, I. Ggsignif: Significance Brackets for ‘Ggplot2’. (2021). |
| library(dplyr) | Wickham, H., François, R., Henry, L., Müller, K. & Vaughan, D. Dplyr: A Grammar of Data Manipulation. (2023). |
| library(survminer) | Kassambara, A., Kosinski, M. & Biecek, P. Survminer: Drawing Survival Curves Using ‘Ggplot2’. (2021). |
| library(gridExtra) | Auguie, B. gridExtra: Miscellaneous Functions for ‘Grid’ Graphics. (2017). |
| library(patchwork) | Pedersen, T. L. Patchwork: The Composer of Plots. (2024). |
| library(ggtext) | Wilke, C. O. & Wiernik, B. M. Ggtext: Improved Text Rendering Support for ‘Ggplot2’. (2022). |
| library(tidyr) | Wickham, H., Davis, V. & Girlich, M. Tidyr: Tidy Messy Data. (2024). |
| library(rstatix) | Kassambara, A. Rstatix: Pipe-Friendly Framework for Basic Statistical Tests. (2023). |
